# Elucidating the Mechanism Underlying UBA7•UBE2L6 Disulfide Complex Formation

**DOI:** 10.1101/2024.11.07.622398

**Authors:** Pei-Tzu Chen, Jia-Yin Yeh, Jui-Hsia Weng, Kuen-Phon Wu

## Abstract

We elucidate cryo-EM structure and formation of the ubiquitin-associated bovine UBA7•UBE2L6 disulfide complex, shedding light on a highly specific and evolutionarily conserved mechanism governing ISG15 conjugation, a pivotal process in the immune response. UBA7 displays a unique capacity to recognize UBE2L6, distinct from this latter’s homolog UBE2L3, highlighting the intricacies of cellular regulation. Inter-species interactions of the resulting complex further underscore its significance. We characterize three crucial factors that influence UBA7•UBE2L6 disulfide complex formation: (1) strong binding affinity and specificity; (2) conformational differences in the catalytic cysteine capping loop (CCL); and (3) increased thiolate/thiol ratios at catalytic cysteines. Modification of any of these factors profoundly impacts complex activation and the ISG15 transfer cascade. This redox-sensitive complex implies a link between oxidative stress and regulation of the immune response, highlighting a potential therapeutic target for modulating immune reactions arising from infections and inflammatory conditions.

## Introduction

Ubiquitin (Ub) and ubiquitin-like (Ubl) proteins—such as ISG15, FAT10, Nedd8, SUMO, and Atg3—share a common structural fold. However, their respective activating enzymes, collectively termed E1, exhibit notable differentiation. For instance, UBA1, UBA6, and UBA7 serve as the E1 enzymes for Ub, FAT10/Ub, and ISG15 activation, respectively. Despite the significant diversity among Ubl proteins, UBA6 is noteworthy for its dual functionality in activating both Ub and FAT10^1,2^. The experimentally determined structures of UBA1^3^, UBA6^4,5^, and UBA7^6^ exhibit striking structural similarities. In contrast, the heterodimeric complexes NAE1•UBA3 and SAE1•SAE2 function as the E1 enzymes responsible for Nedd8 and SUMO activation, respectively. Typically, E1 enzymes comprise multiple domains, including an active adenylation domain (AAD), an inactive adenylation domain (IAD), a first catalytic cysteine half-domain (FCCH), a second catalytic cysteine half-domain (SCCH), and a ubiquitin-fold domain (UFD), with Ubl activation occurring within the AAD, and thioesterification and transthioesterification associated with the FCCH and SCCH, respectively^7^. The C-terminal UFD plays a crucial role in recruiting and recognizing E2 conjugation enzymes^8,9^. Activation of Ub or Ubl proteins by E1 enzymes is initiated upon their consumption of one ATP molecule, resulting in an adenylated Ub or Ubl in the AAD. Subsequently, the Ub or Ubl is transferred to the E1 active site cysteine, forming a thioester bond between the C-terminal glycine carboxyl group and the thiol group of the cysteine. The active site cysteine in E1 is located within the SCCH. Once the E1 is charged with Ub or Ubl, the E2 conjugation enzyme binds to E1 for transthioesterification, transferring Ub or Ubl from E1 to E2 and thereby initiating a subsequent reaction cascade. E2 conjugation enzymes also host an active site cysteine essential for the formation of a thioester product with Ub or Ubl. Forty human E2 enzymes are classified in the InterPro database (https://www.ebi.ac.uk/interpro/), with the majority interacting with UBA1, whereas UBE2L6 and UBE2Z exclusively interact with UBA7 and UBA6, respectively.

It has long been perplexing why E1 and E2 do not spontaneously form a disulfide (SS) complex, given that they both possess individual active site cysteines for transthioesterification, being in close proximity during the reaction. Several factors, including the thiol pKa, entropy, and environmental redox conditions^10^, may play a critical role in preventing the formation of disulfide complexes between these enzymes. However, the precise mechanisms that govern the formation or disruption of such complexes remain unclear. Oxidative conditions have been reported to promote disulfide complex formation for the E1-E2 pairs such as UBA1-Rad6, UBA1-Cdc34, and SUMO E1-Ubc9, indicating a modulated reaction within cells^11–14^. In most cases, the formation of an E1-E2 disulfide complex interrupts Ub (or Ubl) modifications.

Moreover, many structural characterizations of the E1, E2, and Ubl complexes —including *Schizosaccharomyces pombe* Uba1•Ubc15 (PDB ID: 5KNL), *Saccharomyces cerevisiae* Uba1•Cdc34 (PDB ID: 6NYA), and *S. cerevisiae* Uba1•Ubc4 (PDB ID: 4II2)—have been conducted using 2,2’-dipyridyl-disulfide (DSS)^15–17^, which promotes the formation of E1•E2 (• is denoted for the disulfide bond) disulfide complexes in vitro through oxidation.

Immune responses often entail the generation of reactive oxygen species (ROS), antioxidant defense mechanisms, and redox signaling^18^, suggesting that Ub or Ubl enzymes involved in the immune system may be strongly influenced by these environmental changes. Notably, UBA7 is a ubiquitin-modifying enzyme specific to the immune response, where it activates interferon-stimulating gene 15 (ISG15, an Ubl), leading to the ISGylation of numerous cellular proteins^19,20^.

UBE2L6 is known to preferentially transfer ISG15 from UBA7 for subsequent ISGylation^21^. Under conditions of microbial infection, host immune-responsive ROS or redox imbalance may modulate or interfere with the interactions between UBA7 and UBE2L6 during the process of ISGylation.

Here, we reveal that UBA7 and UBE2L6 display a strong preference for forming a disulfide complex *in vitro* without the need for ROS stimulation. We present comprehensive biochemical and structural analyses of the E1 enzymes UBA1 and UBA7, with a focus on their formation of E1•E2 complexes. Utilizing AlphaFold and cryo-EM structures of UBA1, UBA7, and their variants in complex with E2 enzymes, we report the catalytic cysteine capping loop (CCL) as being critical for disulfide complex formation. The distinct affinities for E2 enzymes via the C-terminal UFDs of UBA1 and UBA7 underscore the specificity and catalytic kinetics of transthioesterification, as well as disulfide bond formation. Furthermore, we investigate the pKa values of the active site cysteines of UBA1, UBA7, and their variants, as well as those of the E2 enzymes UBE2L3 and UBE2L6. Our pKa measurements indicate favorable formation of disulfide complex in the presence of low-pKa E2 enzymes without the additional need for ROS such as H_2_O_2_. Collectively, our findings unveil the mechanism underlying spontaneous formation of the UBA7•UBE2L6 SS complex.

## Results

### Formation of a Disulfide-Bonded Complex Between UBA7 and UBE2L6

A critical step in the process of ubiquitination entails the activation and transfer of Ubl to the active site of E2, facilitated by ATP and E1. Following the transfer, E1 can engage in another round of reactions; however, our previous experiments reveal an intriguing anomaly during the UBA7-UBE2L6-ISG15 thioester transfer reaction. Despite adding more ISG15 and UBE2L6, the level of UBE2L6∼ISG15 remained the same. To determine if the transfer efficiency of Ubl proteins from E1 to E2 enzymes can be modulated by altering the order in which E1, E2, and Ubl are mixed, we designed two distinct experimental pathways (**Fig. 1a**). For pathway (i), experimental reactions were initiated by combining E1 enzyme—specifically UBA1 for Ub or bovine UBA7 for ISG15—with ATP and MgCl_2_. Subsequently, we introduced E2 enzymes, i.e., UBE2L3 for Ub or UBE2L6 for ISG15, and allowed the reaction to proceed for 20 minutes.

**Figure 1.**
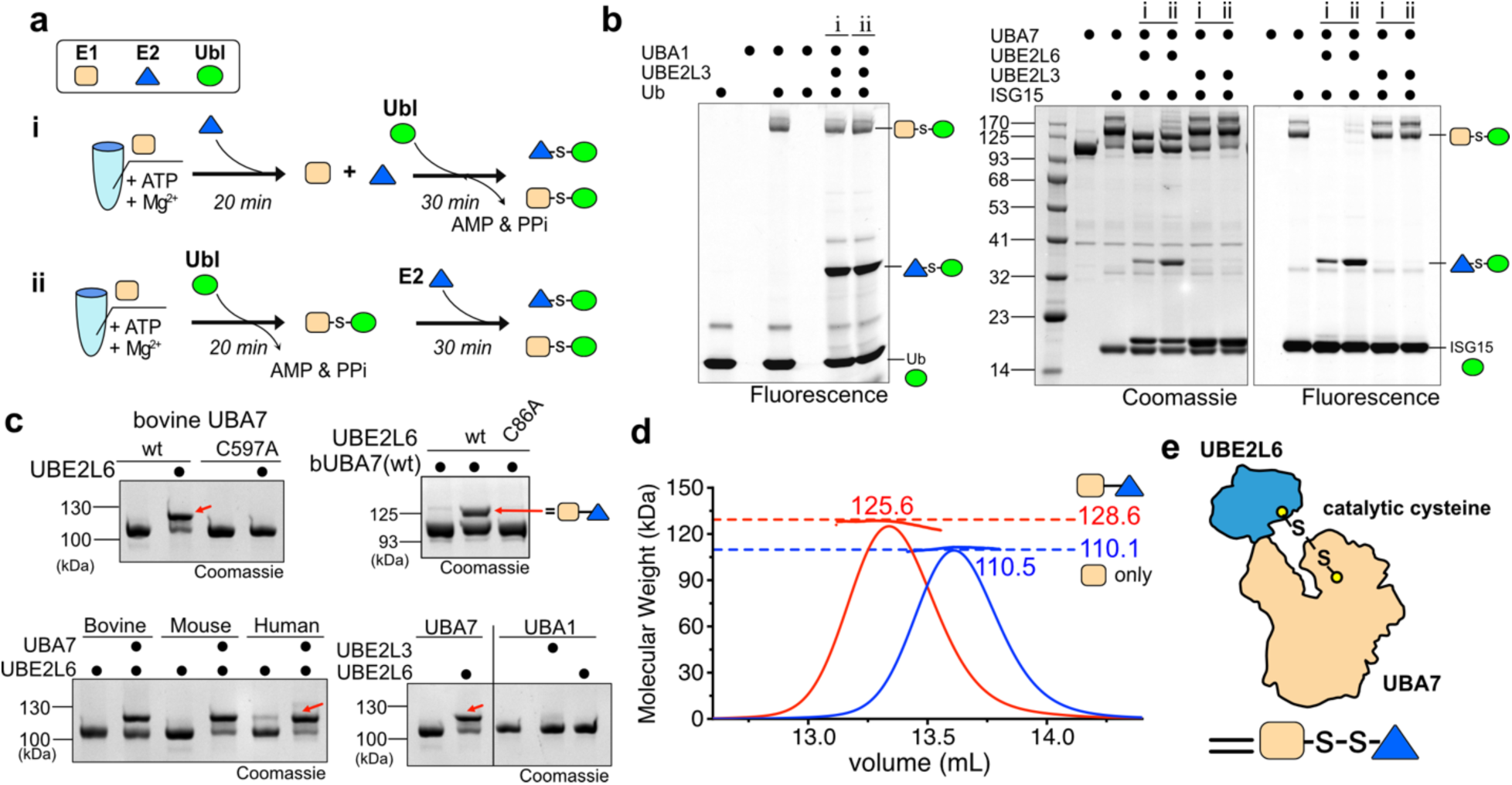
Confirmation of the existence of an E1•E2 disulfide complex. **a.** Two time-dependent chase assays in which the order of addition of an E2 enzyme (UBE2L3 or UBE2L6) or a Ubl (Ub or ISG15) protein is added to the mixture containing an E1 enzyme (UBA1 or UBA7). The expected products are E1∼Ubl and E2∼Ubl if no E1•E2 complex is formed. **b.** The two chase reactions **i** and **ii** produced similar yields of UBA1∼Ub* and UE2L3∼Ub*, where * represents a fluorescein probe. In contrast, the UBA7 plus UBE2L6 premixure in reaction **i** resulted in less UBE2L6∼ISG15* product than detected for the UBA7 plus ISG15* premixure in reaction **ii**. The Coomassie-stained gel shows a high molecular weight (MW) band at ∼125 kDa, which is not ISG15*-related but may be an UBA7•UBE2L6 covalent complex. **c.** The active cysteines in bovine UBA7 (bUBA7) and UBE2L6, respectively, are responsible for UBA7•UBE2L6 complex formation since C597A (UBA7) or C86A (UBE2L6) mutation resulted in no reaction. UBA7-UBE2L6 pairs from human, mouse and bovine all form a disulfide complex (indicated by red arrows), but not human UBA1 mixed with UBE2L3 or UBE2L6. **d.** SEC-MALS chromatographs establish high-precision MWs of UBA7 alone (110.13 kDa theoretical mass) and for the UBA7•UBE2L6 complex. The blue and red dashed lines represent the predicted MWs of UBA7 and UBA7•UBE2L6 complex, respectively. **e.** Cartoon representation of the UBA7•UBE2L6 complex.

**Table 1.** Cryo-EM data collection, refinement, and validation statistics.

|  | UBA7-UBE2L6-ISG15 | MBP-UBA7-UBE2L6 |
| --- | --- | --- |
| Accession code (EMDB) | EMD-61794 | EMD-61979 |
| Accession code (RCSB PDB) | 9JTC | - |
| **Data collection and processing** |  |  |
| Magnification | 130,000 |  |
| Voltage | 300 kV |  |
| Electron exposure (e-/Å<sup>2</sup>) | 50.67 | 50.0 |
| Defocus range (µm) | -1.2 to -2 |  |
| Pixel size (Å) | 0.324 |  |
| Binned Pixel size (Å) | 0.648 |  |
| Symmetry imposed | C1 |  |
| Initial particle images (no.) | 13,956,920 | 5,177,529 |
| Final particle images (no.) | 176,617 | 46,385 |
| Reconstruction method | Single particle reconstruction |  |
| Map resolution (Å) (FSC<sub>0.143</sub>) | 2.99 | 3.83 |
| Map sharpening B factor (Å<sup>2</sup>) | 87.7 | 98.3 |
| **Refinement** |  |  |
| Initial model | AlphaFold 3<sup>45</sup> | - |
| Model composition |  |  |
| Non-hydrogen atoms | 9158 |  |
| Protein residues | 1163 |  |
| Ligand / Water | 1 / 0 |  |
| B factor (Å<sup>2</sup>) (min / max / mean) |  |  |
| Protein | 13.26 / 182.48 / 72.54 |  |
| Ligand | 53.26 / 87.54 / 64.86 |  |
| R.m.s. deviations |  |  |
| Bond lengths (Å) | 0.003 |  |
| Bond angles (°) | 0.541 |  |
| Validation |  |  |
| MolProbity score | 1.19 |  |
| Clash score | 3.62 |  |
| Poor rotamers (%) | 0.91 |  |
| Ramachandran plot |  |  |
| Favored (%) | 97.82 |  |
| Allowed (%) | 2.18 |  |
| Outliers (%) | 0 |  |
| Model vs Data |  |  |
| CC (mask) | 0.72 |  |
| dFSC model (0 / 0.143 / 0.5) (Å) | 1.3 / 1.7 / 3.0 |  |

Next, we introduced Ub/Ubl and continued the reaction for an additional 30 minutes. Alternatively, for pathway (ii), we first mixed Ub/Ubl with E1 enzyme before adding an E2 enzyme into the reaction mixture. As expected, we observed no significant differences in E2∼Ub formation between the two pathways for the UBA1/UBE2L3/Ub system. However, a notable difference was observed in pathway outcomes for the UBA7/UBE2L6/ISG15 system.

Specifically, UBE2L6∼ISG15 formation was less pronounced for pathway (i) compared to pathway (ii), indicating that premixing of E1 and E2 might retard ISG15 transfer **(Fig. 1b)**. Interestingly, in a UBA7/ISG15 charging reaction in which UBE2L6 (the E2 enzyme) was not added, UBA7∼ISG15 formation was clearly detectable, whereas this thioester-linked complex was not detectable in the presence of UBE2L6 **(Fig. 1b)**. We also observed a novel band at ∼125 kDa (*i.e*., between the molecular weights of UBA7 and UBA7∼ISG15) solely in the Coomassie-stained SDS gel of both reaction pathways (i) and (ii).

We hypothesized that UBE2L6 might bind to UBA7 and thereby impede ISG15 loading. Since the putative UBA7•UBE2L6 complex was not disrupted in the SDS gel, we suspected that it formed via a covalent disulfide bond between the two catalytic cysteines of the E1 and E2 enzymes. The addition of β-mercaptoethanol (β-ME) to mixtures containing E1∼ISG15 or E1•E2 complex led to a clear reduction in thioester bonds of UBA7∼ISG15 and UBE2L6∼ISG15* (with * representing a fluorescein probe), as well as the potential disulfide bond of UBA7•UBE2L6 **(Fig. S1**). Given that UBA7 and UBE2L6 could spontaneously assemble and the complex is β-ME-sensitive, we can confirm that the formation of the UBA7•UBE2L6 complex arises via a disulfide bond. This outcome is further validated by the fact that the UBA7•UBE2L6 complex was not detected when either the UBA7(C597A) or UBE2L6(C86A) mutants were employed, underscoring the criticality of the disulfide bond to formation of this complex (**Fig. 1c**, top panels). Intriguingly, we consistently detected UBA7•UBE2L6 complexes in the ISG15-specific human, mouse, and bovine systems, rather than Ub-specific UBA1•UBE2L3 or UBA1•UBE2L6 complexes (**Fig. 1c**, bottom panels). Since proteins of the bovine ISG15 system are well expressed and more stable than those of the human ISG15 system, we focused our biochemical, biophysical, and structural investigations on bovine UBA7, UBE2L6, and ISG15.

To gain a more profound understanding of the molecular mechanism of disulfide complex formation, we measured the molecular weight (MW) of bovine UBA7 (bUBA7) in solution by means of size-exclusion chromatography coupled with multi-angle light scattering (SEC-MALS). The experimentally determined MW of bUBA7 was found to be 110.5 kDa, which closely matches the predicted value of 110.13 kDa. The measured and theoretical MWs of UBA7•UBE2L6 complex are 125.6 kDa and 128.6 kDa, respectively, strongly implying the presence of a 1:1 covalently bonded E1•E2 complex in solution, as illustrated in **Figure 1d-e**. These findings support that bovine UBA7•UBE2L6 is a heterodimeric complex formed by a disulfide bond between the active site cysteines of the two components.

### Structural Analysis of bovine UBA7**•**UBE2L6 and UBA7**•**UBE2L6∼ISG15

UBA7 shares a similar overall architecture with UBA1, as confirmed by previous research^6,22^. However, unlike the highly conserved sequence of UBA1, UBA7 exhibits greater diversity across species. These subtle sequential differences in specific regions may result in different structure different among UBA7 species and play a critical role in the variations observed in its interaction mechanisms. We aimed to determine the cryo-EM structure of bovine UBA7, its covalently bonded partner UBE2L6, and the transient adenylated complex of UBA7•UBE2L6 with ISG15. The structure of bovine UBA7 (bUBA7) may be heterogeneous and dynamic, which complicates the construction of a successful 3D map. To address this, we prepared a maltose-binding protein (MBP)-fused bUBA7, complexed with bUBE2L6 via a disulfide bond formed between the catalytic cysteines of each protein. MBP has previously been used to facilitate protein crystallization in the Nedd8 E1, and similarly, this fusion resulted in a cryo-EM map at 4 Å resolution **(Fig. 2a)**, with clear density for MBP, most regions of bUBA7, and part of bUBE2L6, especially the area near the UFD of bUBA7. We observed that the FCCH and SCCH domains of bUBA7 were relatively dynamic, exhibiting lower local resolution, while the SCCH and UFD domains maintained stable interactions with bUBE2L6. The human UBA7•UBE2L6 structures^6,22^ fit well into the map, except for the more flexible FCCH and SCCH domains. The consistent conformation of the E1-E2 disulfide-bonded complexes suggests that spontaneously formed UBA7•UBE2L6 could represent the ISG15 activation state of UBA7. To further study this, we prepared a mixture of ISG15, ATP, and MgCl₂ with UBA7•UBE2L6, simulating the activation complex of bovine UBA7, UBE2L6, and adenylated ISG15.

**Figure 2.**
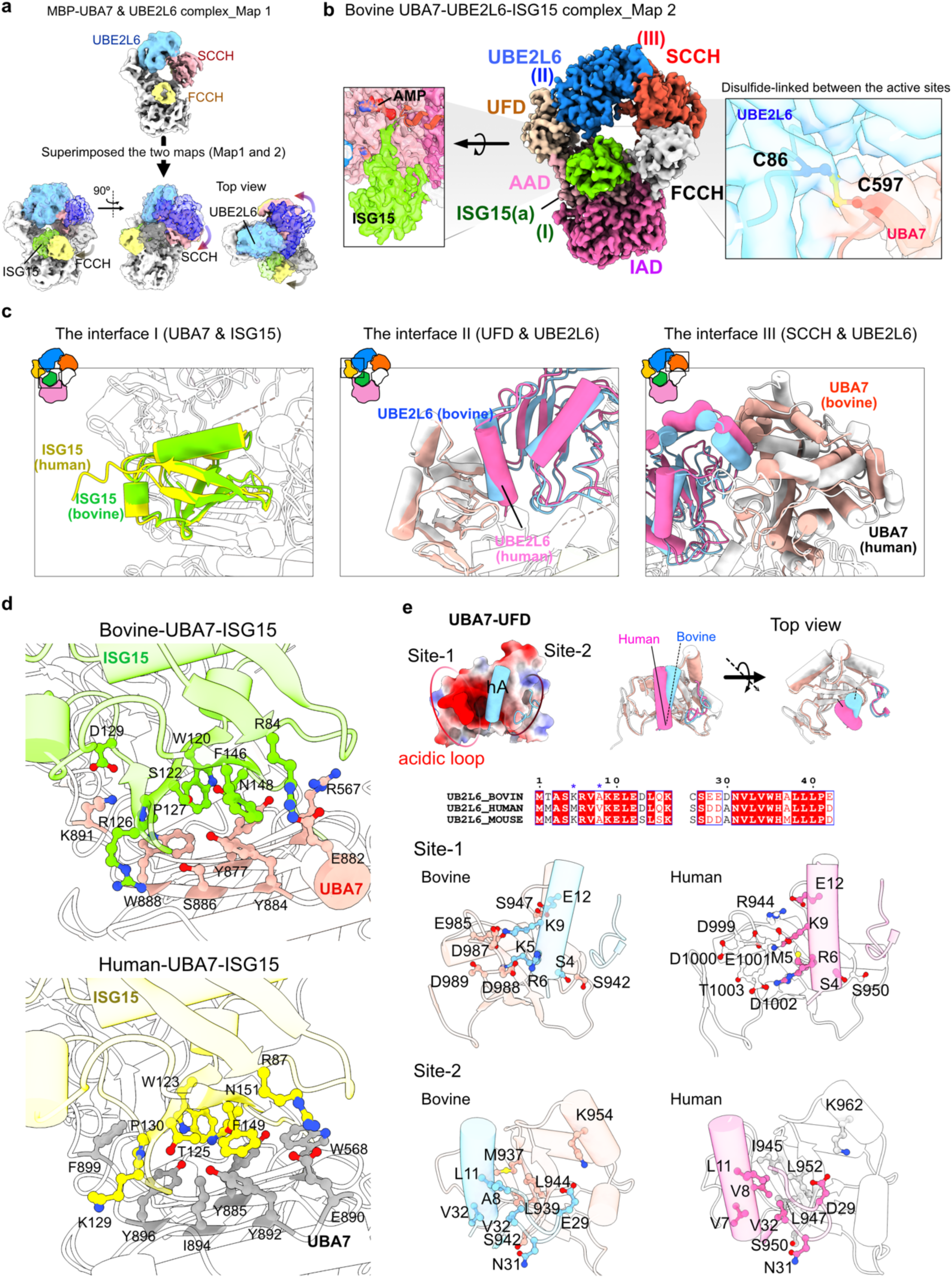
Structural analysis of bUBA7•UBE2L6∼ISG15 complex. **a.** Cryo-EM map of the MBP-UBA7•UBE2L6 complex. UBE2L6 is depicted in sky blue, while the SCCH domain of UBA7 is highlighted in salmon, and the FCCH domain is emphasized in yellow. **b.** Cryo-EM map of the bUBA7•UBE2L6∼ISG15 complex. The domains of UBA7 are displayed in the same color codes, while ISG15 is represented in green. Zoomed-in views show the cryo-EM density of adenylated ISG15 bound to UBA7, as well as the disulfide bond formed between the catalytic cysteines of UBA7 and UBE2L6. **c.** Superimposed structures of three key interfaces within the UBA7/UBE2L6/ISG15 complex from both bovine and human (PDB ID: 8SEB). The interfaces include: (I) UBA7/ISG15, (II) UBA7(UFD)/UBE2L6, and (III) UBA7(SCCH)/UBE2L6. **d.** Comparative analysis of the AAD and ISG15 interaction in the UBA7/UBE2L6/ISG15 complex between bovine and human. Structural alignment highlights the interaction of the AAD of UBA7 with ISG15, revealing both conserved and distinct interaction patterns across species. Key contact residues are shown, emphasized by the ball and stick style. **e**. Interface analysis of the UFD and E2 structures in the UBA7/UBE2L6 complex. The UFD of bovine UBA7 is displayed as a colored surface representing electrostatic potential, while the hA helix of UBE2L6 is shown as a cartoon. Based on electrostatics, the left and right areas of the hA are identified as interaction Site-1 and Site-2, respectively. The superimposition of the bovine and human’s UFD of UBA7 (bovine’s colored in light salmon and human is white) and UBE2L6 (bovine: sky blue, human: pink) structure, illustrates structural differences. Sequence alignment of the different species’ UBE2L6. The blue asterisk marked the residues that differ between the UBE2L6s’ hA of bovine and human. Magnified views of Site-1 and Site-2. The structures are represented as transparent cartoons and the residues contained in the contact are shown as ball and stick.

The cryo-EM map of the activation complex, reconstructed from 18,731 images, achieved a resolution of 2.99 Å **(Fig. 2b)**, clearly resolving the three component proteins. The adenylated C-terminal region of bISG15 is well-defined in the map, while the N-terminal domain of bISG15 (residues 1-79) remains dynamic and is not visible. Bovine UBE2L6 tightly interacts with the UFD and SCCH domains of bUBA7, and the disulfide bond between C597 of bUBA7 and C86 of bUBE2L6 is clearly observed **(Fig. 2b)**. The bovine ISG15 activation complex aligns well with the human ISG15 activation complex^6^ (PDB ID: 8SEB), with an RMSD of 1.27 Å across 59,616 aligned atoms. For example, the C-terminal lobes of ISG15 **(Fig. 2c)**, and the IAD, AAD, and UFD domains of UBA7 from both species, are perfectly aligned. However, in our bovine ISG15 activation complex compared to the MBP-UBA7•UBE2L6 complex, we observe a slight displacement of UBE2L6 and an upward lift of the SCCH domain of UBA7, even though the structures are disulfide-bonded between UBE2L6 and UBA7.

The aromatic hydrophobic interactions between the C-lobes of bovine and human ISG15 and their respective UBA7 partners are conserved. For example, W120 and F146 of the bISG15 C-lobe make close contact with Y877, Y884, and W888 of bUBA7. This interaction is similarly observed in human ISG15 (W123 and Y149) with hUBA7 (Y885, Y892, and Y896) **(Fig. 2d)**. Additionally, a salt bridge is formed between R84 of ISG15 and R567 of UBA7, along with D129-K891, and 17 hydrogen bonds, including R84-E882, R126-R887, and P127-K891, which stabilize adenylated ISG15 on bovine UBA7. Similar interactions were observed in the human ISG15 activation complexes (PDB IDs: 8OIF and 8SEB).

UBE2L6 primarily interacts with UBA7 via the SCCH and UFD domains. In the bovine ISG15 activation complex we determined, the first helix (hA) of UBE2L6 is positioned at the interface of an acidic patch (contact site I) and a relatively neutral region (contact site II) on the UFD domain **(Fig. 2e)**. Interestingly, the orientation of the hA helix in bovine UBE2L6 is shifted toward site II, while in the human complex, it is directed toward site I, resulting in a 20° rotation. For instance, residues K5 (M5 in human), K9, and E12 in bovine UBE2L6 form salt bridges and hydrogen bonds with site II of the UFD. Meanwhile, A8 (V8 in human), L11, and V32 of UBE2L6 make hydrophobic contacts with L939 and L944 (L947 and L952 in human) of the UFD. The cryo-EM structures characterized the distinct UBE2L6-UBA7 interactions between human and bovine species, highlighting significant specificity. Validation of the cross-species E1-E2 disulfide bond complex further confirmed the interaction barrier, as demonstrated in the subsequent biochemical reaction analysis. **(Fig. S5)**.

### Structural Analysis of the Catalytic Cysteine Capping Loop (CCL) in UBA7**•**UBE2L6

UBE2L6 also interacts with the SCCH domain in UBA7. In the bovine UBA7•UBE2L6∼ISG15 complex, frequent hydrogen bonds were identified between UBE2L6 and the CCL of UBA7, including interactions such as D80 (UBE2L6)-R600 (UBA7), R81-Q645, R81-D774, and R122-D774 near the disulfide-bonded catalytic cysteines, and E124-S769, E127-R688, and T130-K694 locate on the helix C and packs against the SCCH domain. The human UBA7 activation complex displays similar interactions between hUBE2L6 and the catalytic center in the SCCH domain.

Both our bovine and previously published human UBA7 activation complex structures revealed that the CCL is distant from the catalytic cysteines, C597 in bUBA7 and C599 in hUBA7 **(Fig. 3a, b)**. The CCL in bUBA7 is a short loop near the catalytic cysteine, with its boundaries defined by the N-terminal (entry) and C-terminal (exit) helices. We further analyzed the lengths of these CCLs by structurally aligning *S. cerevisiae* UBA1 (yUBA1, PDB ID: 7ZH9), human UBA7 (PDB ID: 8SEB), and bUBA7 (this study).

**Figure 3.**
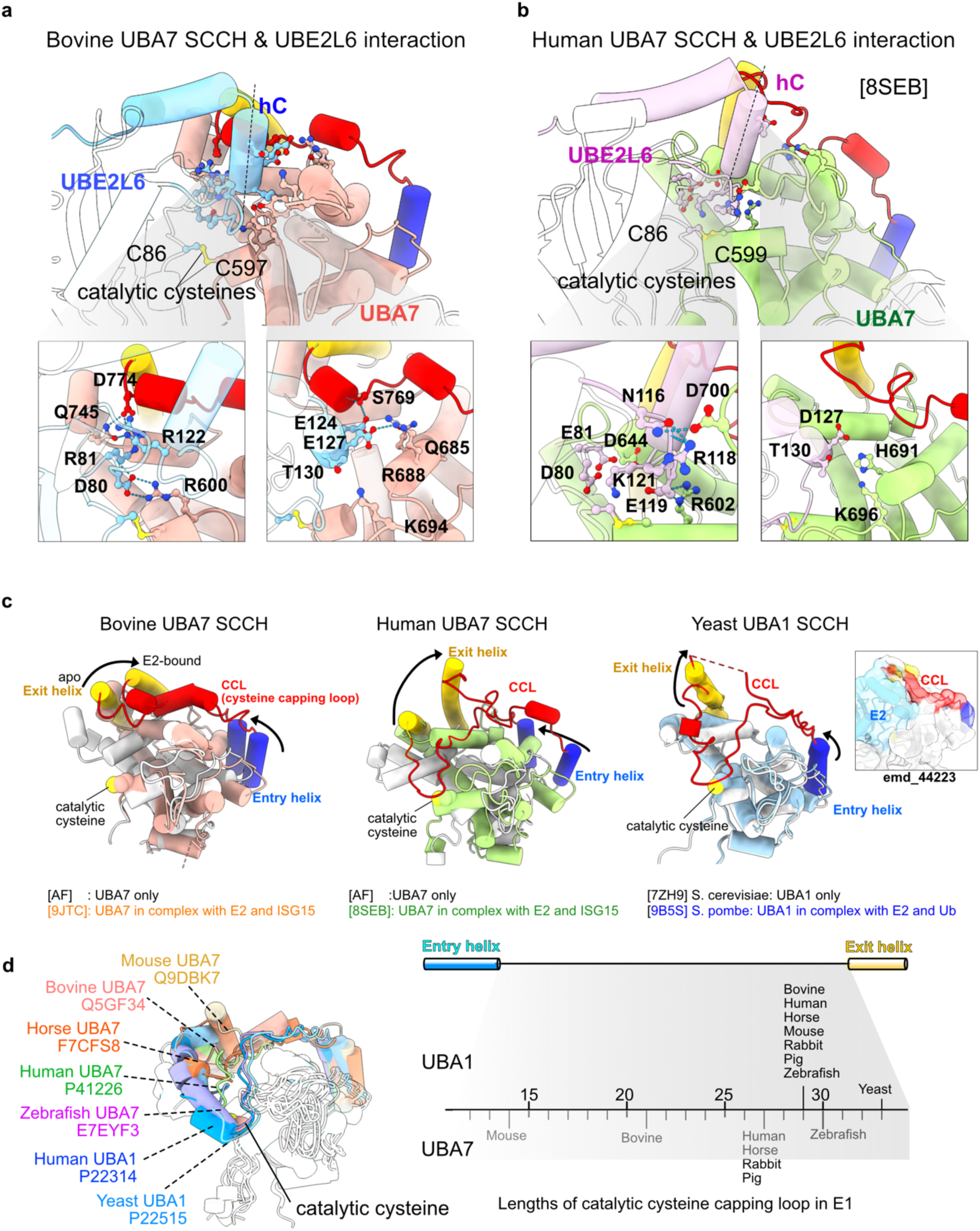
The role of the SCCH domain in the UBA7 and UBE2L6 interaction. **a, b.** Magnified views of the interface between the SCCH domain of UBA7 and UBE2L6 in both bovine and human structures. Contacting residues at the interface are depicted in ball-and-stick representation. **c.** Structural alignments of the SCCH domain of E1 in its apo form and in complex with E2. The catalytic cysteine capping loop (CCL) is defined by the loop structure flanked by two helices: an entry helix at the N-terminal and an exit helix at the C-terminal ends. In the yeast E1-E2 complex model, the CCL region is partially missing, but it becomes visible in the electron density map (EMD_44223) when the contour level is lowered. **d.** Superimposed structural models of the SCCH domain from UBA7 and UBA1 across different species, obtained from the AlphaFold database (UniProt IDs displayed below each species’ name). The entry and exit helices, as well as the catalytic cysteine capping loop (CCL), are highlighted in color, while the rest of the structure remains transparent. The number of residues in the CCL for each species is counted and annotated along a number line.

The CCL is positioned away from the catalytic cysteines in both the human and bovine UBA7 activation complex structures. However, the behavior of the CCL in the apo form of UBA7 remains undetermined, as it has not yet been experimentally characterized. To explore this, we downloaded free forms hUBA7 and bUBA7 from the AlphaFold protein structure database (https://alphafold.ebi.ac.uk/). Superimposing the free and E2-disulfide-bonded forms of bUBA7, hUBA7, and yUBA1 revealed conformational changes in the SCCH domain. The exit helix consistently extends outward from the catalytic cysteine when covalently bonded to E2. In the AlphaFold model of free bUBA7, closer examination of the SCCH domain, along with yUBA1, revealed that C597 of bUBA7 is highly exposed, while C600 of yUBA1 is shielded by a flexible catalytic cysteine capping loop (CCL).

In contrast to the indirect contact between the CCL and the catalytic cysteine in bUBA7 and hUBA7, the CCL in yUBA1 covers the catalytic cysteine in the apo form and flips over upon loading of the E2 enzyme^23^(PDB ID: 9B5S) **(Fig. 3c)**. This suggests a potential mechanism in UBA1 that long CCL might prevent rapid disulfide bond formation with multiple E2 enzymes. To further investigate this, we downloaded AlphaFold models of the apo forms of seven UBA7 proteins and compared them to the free forms of hUBA1 and yUBA1. The exit helix and CCL displayed significant conformational diversity across the nine structures **(Fig. 3d)**, with the CCLs in hUBA1 and yUBA1 fully sheltering the catalytic cysteine. The positioning of the exit helix in UBA7 varies considerably across species, reflecting the variability in CCL length, which ranges from 14 residues in mouse to 21 in bovine, 27 in human and horse, and 30 in zebrafish compared to long CCL in most UBA1 (29 residues).

### Dynamic Nature of the SCCH Domain

Cryo-EM maps of MBP-fused bUBA7•UBE2L6, bUBA7•UBE2L6∼ISG15, hUBA7•UBE2L6∼ISG15, and doubly ISG15 loaded UBA7•UBE2L6 consistently presented weak and relative low-resolution SCCH domain (**Fig. S2, S3**). Such weak SCCH signals in cryo-EM maps imply that the SCCH domain undergoes dynamic conformational changes in solution. To further assess the dynamic nature of the SCCH and the entire UBA7 enzyme, we employed hydrogen-deuterium exchange mass spectrometry (HDX-MS), a powerful technique for rapidly assessing protein structure and dynamics^24^, allowing us to compare structural differences between UBA7 and UBA1, as well as conformational changes upon binding E2 enzymes **(Fig. 4)**.

**Figure 4.**
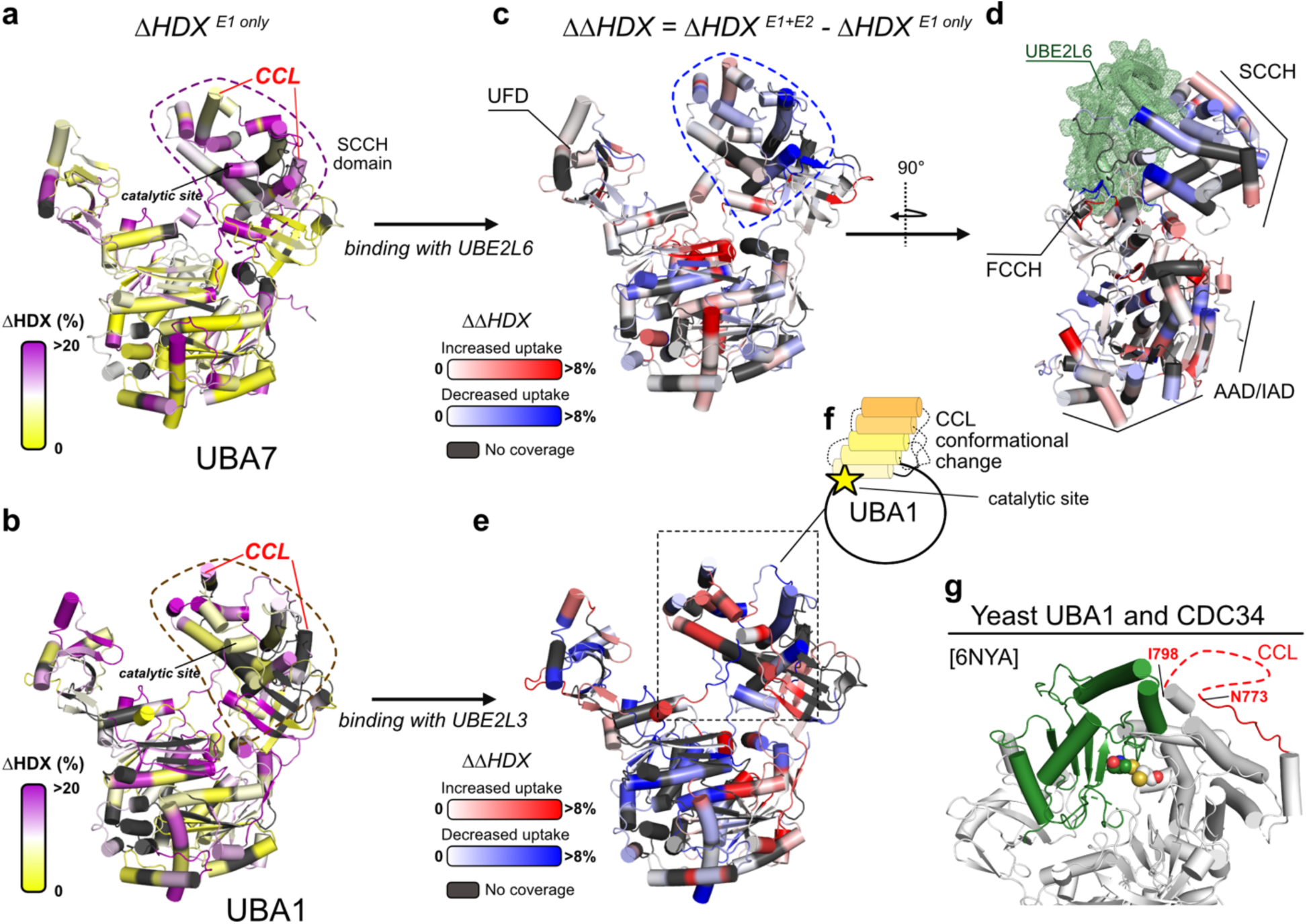
HDX-MS reveals dynamic structural characteristics of E1 enzymes. **a-b.** Bovine UBA7 and human UBA1 display structural similarities, yet the hydrogen-deuterium exchange rates in their respective SCCH domains are substantially different. The UBA7 SCCH domain exhibits a mean ΔHDX >15%, whereas ΔHDX values for most of the residues in the UBA1 SCCH are <5%. **c-d.** Upon UBA7 binding to UBE2L6, ΔΔHDX values for the UBA7 SCCH are notably reduced, implying that UBE2L6 stabilizes SCCH. **e.** When human UBA1 binds UBE2L3, ΔΔHDX values for the UBA1 SCCH region are notably increased. In this scenario, the CCL is more exposed to the solvent (resulting in increased ΔΔHDX), implying the CCL is in an open state (as illustrated in **f**)**. g.** The dynamic characteristics of the CCL of UBA1 are consistent with the crystal structure of the yeast UBA1•CDC34 disulfide complex (PDB ID 6NYA), for which the structure of the CCL spanning N773-I798 is too dynamic to be resolved.

The AAD and IAD domains of bUBA7 alone were well protected from solvent exposure according to HDX-MS, resulting in low deuterium incorporation (ΔHDX <5%). In contrast, the SCCH domain, and particularly the CCL region, exhibited relatively higher dynamics, with 10-20% of amide protons undergoing deuterium exchange (ΔHDX 10-20%). Thus, our HDX-MS analysis supports the dynamic nature of the SCCH domain of UBA7 **(Fig. 4a)**, consistent with our and previous cryo-EM works^6,22^. In contrast, UBA1 alone presented modest levels of deuterium incorporation throughout. Indeed, the ΔHDX value for the entire SCCH domain of UBA1 was not significantly higher than for other regions of UBA1 **(Fig. 4b)**. These results imply distinct intrinsic dynamics between UBA7 and UBA1, which could be related to how they interact with E2 enzymes.

Given that structural changes of E1 and E1•E2, differences in deuterium incorporation between the *apo* and E2-bound states of E1 enzymes would likely illuminate structural changes in these latter. E1•E2 complex formation is initiated by interactions between E2 and the UFD domain of E1^15–17^, followed by Ub conjugation to E2 at the catalytic cysteine in its SCCH domain. Upon binding of bUBE2L6, we detected no significant changes in deuterium uptake in the UFD of bUBA7, although some regions in the UFD were not detected by HDX-MS. However, bUBE2L6 binding did significantly reduce deuterium incorporation by the catalytic patch of the SCCH domain of UBA7, which includes its CCL **(Fig. 4c-d)**. The enhanced stability of bUBA7’s SCCH is consistent to our obtained cryo-EM maps. Moreover, in response to UBE2L6 binding, the AAD and IAD domains of UBA7 underwent local conformational changes, resulting in increased or decreased deuterium uptake. hUBA1 displayed more pronounced changes relative to bUBA7 in deuterium incorporation upon hUBE2L3 binding, with substantially reduced deuterium exchange in the E2-bound state, resulting in a more stable AAD/IAD region than the corresponding region in bUBA7. Remarkably, the CCL of the SCCH domain in hUBA1 exhibited greater deuterium uptake upon interaction with UBE2L3, indicating increased CCL dynamics. This observation aligns with the elevated intrinsic dynamics of the CCL of UBA1 observed in the presence of an E2 enzyme, as confirmed by the crystallographic structures of UBA1•CDC34 **(Fig. 4g)** (PDB ID: 6NYA) ^16^. These findings indicate that E2 binding induces conformational changes in the CCL of the E1 binding partner (**Fig. 4f**).

### Role of the CCL and UFD in Transthioesterification

Given the critical role of the CCL in driving or preventing formation of the E1•E2 (SS) complex, we hypothesized that generating chimeric E1 variants with CCLs of altered lengths would impact disulfide bond formation. Accordingly, we created two E1 variants by either shortening or extending the length of the CCL in UBA1 or UBA7, respectively, resulting in UBA1(dCCL) (where dCCL represents deletions in the CCL) and UBA7(eCCL) (where eCCL represents extensions in the CCL). Additionally, we swapped their UFD domains to generate UBA1(U7) (U7: UFD of UBA7) and UBA7(U1) (U1: UFD of UBA1) chimeras **(Fig. 5a, Fig. S4)**. The AlphaFold-generated model of UBA7(eCCL) depicted the extended CCL as mimicking the UBA1 CCL in that it moved closer to catalytic cysteine C597, resulting in a less solvent-exposed cysteine. Conversely, the AlphaFold-generated model of UBA1(dCCL) showed an unprotected C632, which could potentially affect E1-E2 interactions or enzymatic catalysis.

**Figure 5.**
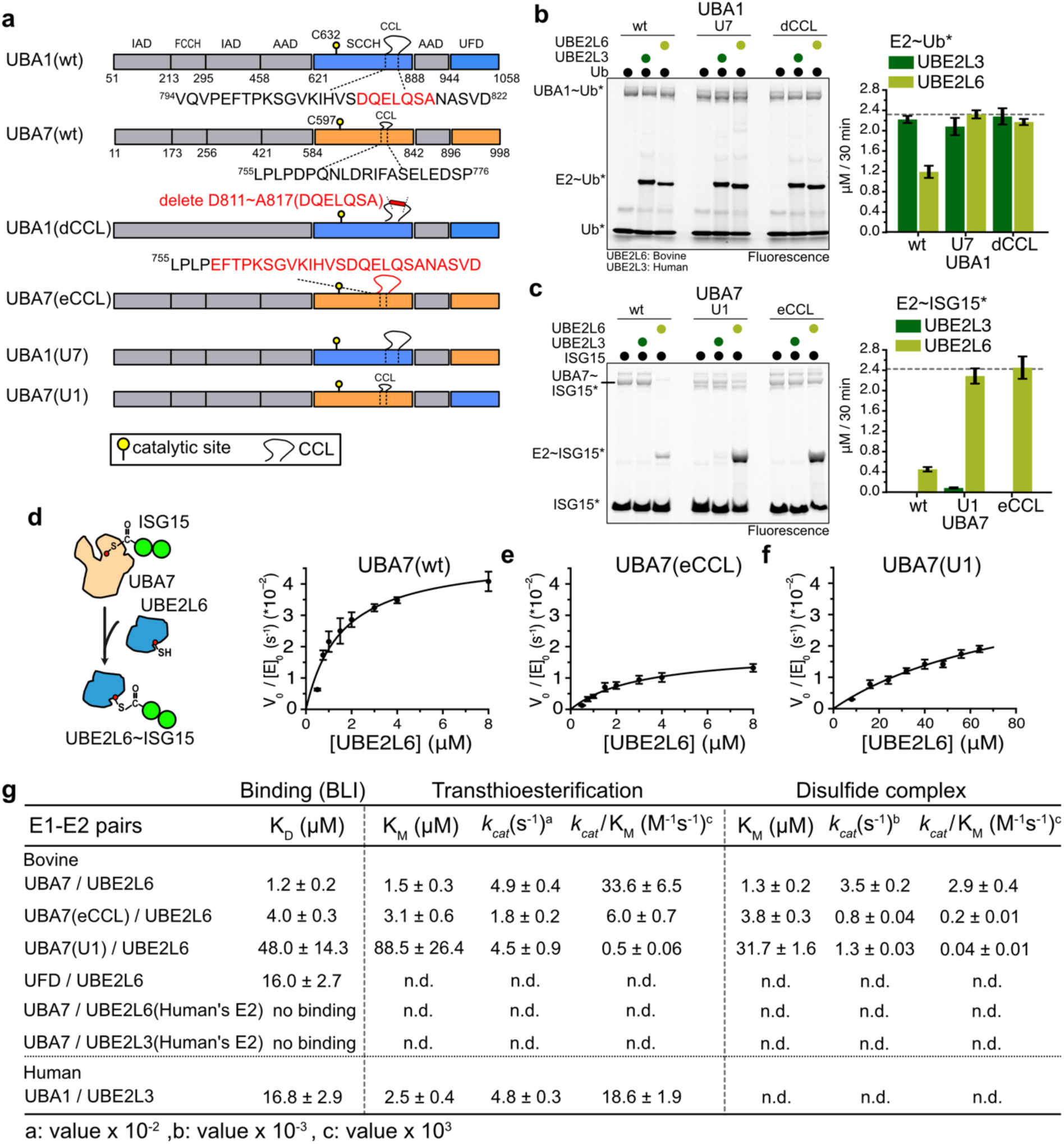
The enzymatic kinetics of UBA1, UBA7 and their variants. **a.** Schematics of the domains in UBA1 and UBA7, as well as their variants. The deleted and extended sequences in UBA1(dCCL) and UBA7(eCCL) are indicated. Chimeras in which the UFD was swapped between UBA1 and UBA7 are also shown. In the chimeras, the CCL remains unchanged. **b-c.** Production yields of E2∼Ub* or E2∼ISG15* (E2 = human UBE2L3 or bovine UBE2L6) using a wildtype (wt) E1 enzyme (UBA1 or UBA7) or a mutant variant. The reaction comprised a mixture of E2 enzyme, Ubl (Ub or ISG15), MgCl2 and ATP, and it was initialized by adding an E1 enzyme into the solution. The 30-min reaction was evaluated by SDS-PAGE. For hUBA1, both UBE2L3 and UBE2L6 can transfer Ub* from wt UBA1. UBA1(dCCL) and UFD-swapped UBA1(U7) generated elevated levels of UBE2L6∼Ub*, but they had no significant impact on UBE2L3∼Ub* production. Note, UBA7 exclusively binds to UBE2L6 as its cognate E2 enzyme for ISG15* transfer. When UBA7(eCCL) or UFD-swapped UBA7(U1) were used as the E1 enzyme, UBE2L6∼ISG15* production was dramatically increased compared to when wt UBA7 was used. UBA7(U1) could also conduct a small amount of UBE2L3 transthioesterification to generate UBE2L3∼ISG15*. **d-g.** The Michalis-Menten kinetics of wt UBA7, UBA7(U1), and UBA7(eCCL), revealing diminished kcat values for the mutant variants. The dissociation constants (KD, Bio-Layer Interferometry, BLI) and Michalis-Menten constants (KM) calculated for wt UBA7, UBA7(eCCL), and UBA7(U1) are highly consistent. For example, the UBA7(U1) chimera exhibits reduced KM and KD values (50-90 µM) relative to wt UBA7 **(g)**. The E1 or E2 enzymes used in the BLI assay are catalytic dead to prevent the spontaneous disulfide complex formation.

To validate these findings, we conducted assays to assess the formation of charged E1∼Ubl and the transthioesterification product E2∼Ubl **(Fig. 5b-c)**. There were no significant differences in charging reactions of UBA1∼Ub among the three UBA1 variants, confirming that neither dCCL nor the swapped UFD greatly impacted Ub thioesterification (**Fig. 5b**). However, the efficiency of Ub transthioesterification from E1 to E2 varied between UBE2L3 (human) and UBE2L6 (bovine) when wildtype (wt) UBA1 was used. The UBA1(U7) chimera displayed a ∼2-fold increase in UBE2L6∼Ub formation, similar to the level of accumulation determined for UBE2L3∼Ub. Shortening CCL length in UBA1 resulted in increased levels of UBE2L6∼Ub, whereas levels of UBE2L3∼Ub were unchanged. The UBA7 variants displayed similar amounts of UBA7∼ISG15 product, consistent with those of the UBA1 variants **(Fig. 5c)**. UFD-swapped UBA7(U1) marginally facilitated UBE2L3∼ISG15 formation, but it dramatically increased (6-fold) UBE2L6∼ISG15 formation in 30 minutes. Compared to wt UBA7, for which UBE2L6 and ISG15 compete for disulfide bond formation and charging, the extended CCL variant may prevent UBE2L6 from forming disulfide bonds, allowing UBE2L6∼ISG15 to accumulate. These results support that changes in UFD and the CCL of E1 enzymes indeed alter E1 catalysis.

To further investigate the impact of the CCL and UFD on E1 catalysis, we measured the enzymatic kinetics of UBA7 variants (**Fig. 5d-f**). For transthioesterification using UBE2L6 (UBE2L6∼ISG15), UBA7 exhibited a kinetic turnover efficiency (*k_cat_*/K_M_) of 33.6 x 10^3^ M^-1^s^-1^. Extending the CCL (eCCL) of UBA7 reduced its *k_cat_*/K_M_ value to one-fifth that of wt UBA7, with a 2.7-fold decrease in both the *k_cat_* and a 2-fold increase in K_M_ values. UFD-swapped UBA7(U1) was profoundly inactive, with a *k_cat_*/K_M_ value 60-fold lower than that of wt UBA7, mainly due to a significantly increased K_M_ value between UBA7 and UBE2L6. Interestingly, UBA7(eCCL) and UBA7(U1) were less catalytically efficient than wt UBA7. Next, we determined the binding affinity between UBA7(C597A was used to avoid disulfide bond formation) and UBE2L6 by means of Bio-Layer Interferometry (BLI), which revealed a dissociation constant (K_D_) of 1.2 µM, reflecting a strong affinity (**Fig. 5g, Fig. S5**). This affinity was slightly reduced to 4.0 µM when we used the extended CCL variant UBA7(eCCL). These binding constants align with the Michaelis-Menten constants (K_M_) shown in **Figure 5g**. In contrast, UFD-swapped UBA7(U1) and wt UBA1 displayed weak K_D_ values of 48.0 µM and 16.8 µM for binding to UBE2L6 and UBE2L3, respectively. The UBA7-UBE2L6 complex exhibited a ∼15-fold tighter binding affinity than the UBA1-UBE2L3 complex, reflecting a readily formation of the disulfide-bonded UBA7•UBE2L6 complex.

Next, we established the formation kinetics of the E1•E2 (SS) complex (**Fig. 6a-d**). We calculated a *k_cat_*/K_M_ value of 2.89 x 10^3^ M^-1^s^-1^ for the wt bUBA7•UBE2L6 complex, with a K_M_ of 1.3 µM consistent with its K_D_ value (1.2 µM). Extending CCL reduced the *k_cat_*/K_M_ value by a factor of 15 for the UBA7(eCCL)•UBE2L6 complex. In line with E1-E2 binding interactions, we detected a substantial decrease in disulfide-bonded complex formation upon using UBA7(U1), with catalytic turnover of UBA7(U1) declining 70-fold relative to wt UBA7. Although these variants exhibited reduced activity, UBA7(eCCL) inhibited formation of disulfide-bonded E1•E2 complexes, allowing for multiple rounds of transthioesterification between UBA7 and UBE2L6∼ISG15. Collectively, these results highlight the roles of the CCL and UFD in modulating E1 catalysis and the formation of covalent E1•E2 complexes.

**Figure 6.**
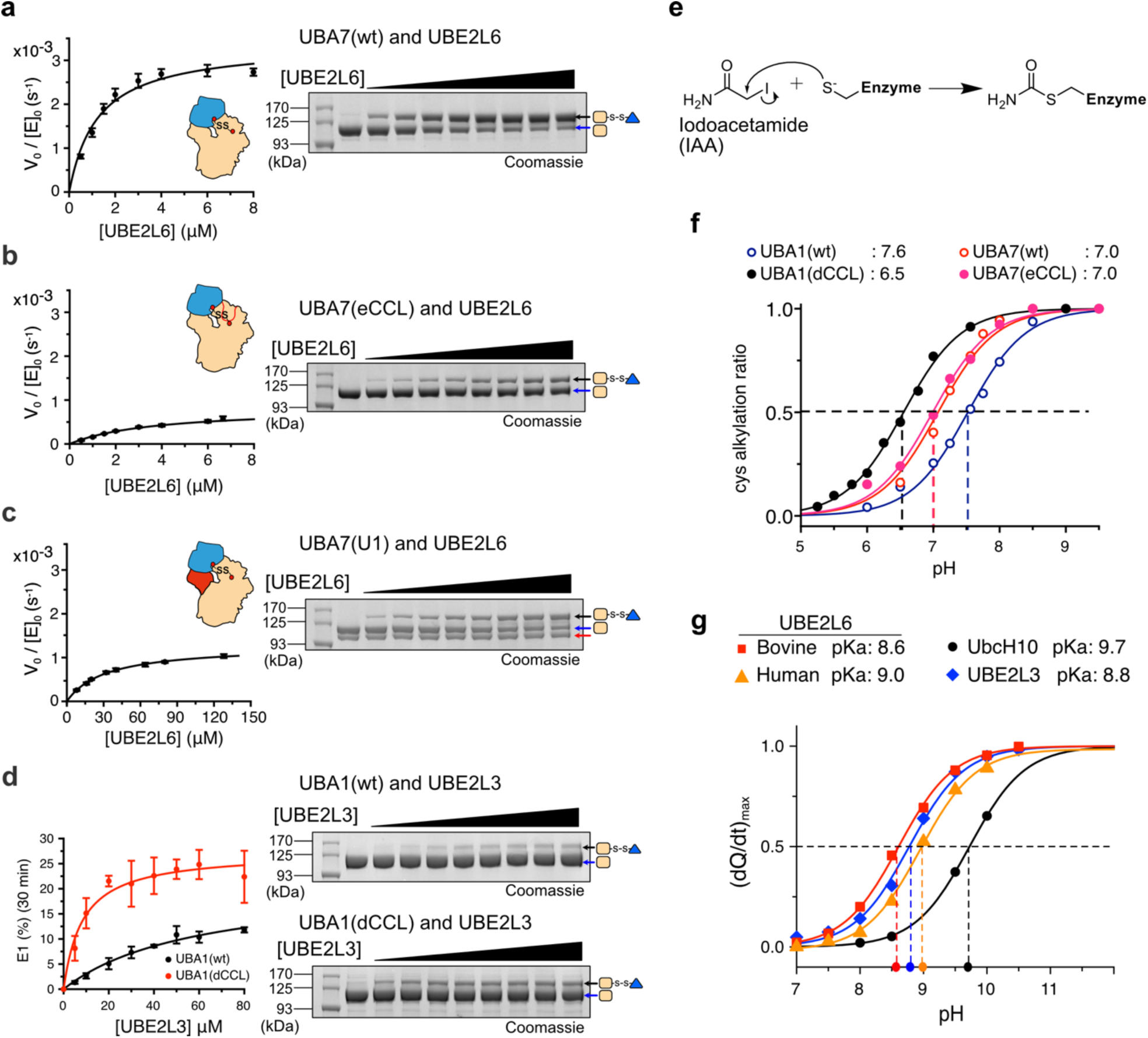
The disulfide complexes of UBA7 variants and the pKa scores of selected E1-E2 pairs. **a-c.** The UBA7•UBE2L6 complex was monitored using SDS-PAGE by titrating UBA7 with varied concentrations of UBE2L6. Production of the complex was assessed and the data was used to fit the kinetics profile. Production of the complex was significantly reduced upon using the UBA7(eCCL) variant, as its extended CCL covers the catalytic cysteine. The UFD-swapped UBA7(U7) variant presented both reduced catalytic efficiency and binding affinity. The black and blue arrows in the SDS gels indicate bands for the disulfide complex and the E1 enzyme alone, respectively. The red arrow in **c** denotes UBA7(U1) variant contamination. **d.** Evaluation of disulfide complex formation for hUBA1 and the UBA1(dCCL) variant with UBE2L3. Due to low binding affinity and catalytic efficiency, only a single reaction at 30-min was collected. The ratio of disulfide complex production over the total amount of E1 enzyme was calculated. The hUBA1(dCCL) variant significantly enhances disulfide complex formation. **e.** Schematic of the alkylation reaction, using iodoacetamide to react with the thiolate group of cysteine. **f.** The alkylation ratios of UBA1, UBA7, and their variants are pH-dependent. **g.** The alkylation ratios of various E2 enzymes (single cysteine variants) were directly measured according to changes in heat release in different pH buffers. In **f** and **g**, datapoints for the E1 and E2 enzymes were fitted to determine pKa values.

In contrast, we hardly detected the formation of UBA1•UBE2L3 complex. In fact, the formation rate was too low to precisely measure the *k_cat_* and K_M_ values, regardless of whether wt UBA1 or UBA1(dCCL) was used (**Fig. 6d**). However, UBA1(dCCL) did facilitate formation of disulfide-bonded E1•E2 complexes (Fig. 6d). AlphaFold-based predictions of complex structures using UBA1(dCCL) and UBE2L3 further confirmed the potential for a disulfide-bonded complex (**Fig. S6**). These data echo the reduced formation of E1•E2 complexes upon use of UBA7(eCCL) (**Fig. 6b**).

### Impact of Cysteine Thiol Deprotonation

In addition to our assessments of the CCL and UFD of E1 enzymes, we also explored the sidechain redox states (protonated: SH, deprotonated: S^-^) of catalytic cysteines in both E1 and E2. The balance of thiol/thiolate groups is affected by the acid dissociation constant (pKa) of sulfenic acids, with the more thiolate (S^-^) cysteines present, the greater the propensity for stimulating disulfide formation. We measured the pKa of E1 and E2 enzymes by mixing them with cysteine-reactive iodoacetamide^25–27^ under varied pH conditions and then monitoring thioester reactivity or released heat, respectively (**Fig. 6e**).

We found that E1 enzymes were favorably alkylated by iodoacetamide at basic pH, resulting in almost no free E1 available for E1∼Ubl thioester formation. Conversely, at acidic pH, the E1 enzymes remained unalkylated, leading to abundant E1∼Ubl thioester formation **(Fig. S6)**. Next, we determined pKa values from the pH-dependent curves of alkylated UBA1, UBA7, and their CCL variants (**Fig. 6f**). Both wt UBA7 and UBA7(eCCL) displayed a pKa of 7.0, whereas the pKa for UBA1(dCCL) was significantly reduced by one pK unit relative to wt UBA7. This lower pKa value for UBA1(dCCL), indicative of increased thiolate at C632, could explain the enhanced reactivity of this variant in terms of disulfide bond formation, as confirmed in **Figure 5d**.

We also directly monitored changes in heat signatures of E2 catalytic cysteines using a method reported previously^25^. Non-catalytic cysteines were mutated to serine in all E2 enzymes to avoid changes in heat arising from undesirable alkylation. Previously reported high pKa values, ranging from 10.2 to 11.1, for the active site cysteines of UbcH10, Ubc2, and Ubc13 imply general activity in preventing E2 enzymes from rapidly reacting with environmental electrons^26^. However, we determined pKa values in the range of 8.6-9.0 for UBE2L3 and UBE2L6 (both human and bovine), *i.e*., 1 to 1.5 pK units lower than reportedly high-pKa E2 enzymes. Without a surprise, UBE2L3 is evolutionally close to UBE2L6^28^ resulted in similar pKa values. For bovine UBE2L6, with a pKa of 8.6, the ratio of thiolate/thiol of the catalytic cysteine at pH 8.0 is 0.25, indicating that 20% of the active site cysteines are readily reactive for disulfide bond formation (**Fig. 6g**), implying potential UBA7•UBE2L6 complex formation at pH 8.0.

Given that the pKa values of E1 and the CCL variants ranged from 6.5-7.6, the catalytic cysteines of UBA1 and UBA7 mainly exist in the thiolate form, making them readily available for Ubl thioesterification and subsequent cascade reactions. The low pKa values of UBE2L6 and UBE2L3 result in the presence of thiolate cysteines in the E2 enzymes, thereby facilitating instantaneous formation of disulfide complexes. UBE2L3 natively doesn’t form disulfide complex with UBA1, however, the decreased CCL UBA1 variant no longer prevents UBE2L6 from direct cysteine-cysteine contacts. Conversely, UbcH10 exhibits a high pKa value of 9.7, with only 1% of thiolate cysteine occurring at pH 8.0 and thus preventing this E2 enzyme from readily forming disulfide bonds (**Fig. 6g**).

## Discussion

In this study, we have comprehensively characterized the intricate interplay between UBA7, an E1 enzyme specific to ISG15, and its cognate E2 enzyme, UBE2L6. Our findings illuminate how a unique and covalent disulfide complex forms between UBA7 and UBE2L6, which is conserved across humans, mice, and bovines. This phenomenon diverges significantly from the conventional non-covalent interactions observed for other E1-E2 pairings. Our discovery provides profound insights into the regulatory mechanisms underlying ISG15 conjugation and the broader realm of ubiquitin-like modifications.

We commenced our analyses by dissecting the remarkable specificity exhibited by UBA7 for UBE2L6 over its homologous counterpart UBE2L3. Despite considerable sequence identity between UBE2L6 and UBE2L3 (55 %), UBA7 exhibits a clear preference for forming disulfide complexes with UBE2L6. This specificity remains consistent even when cross-species interactions were explored, further underscoring the unique nature of the UBA7-UBE2L6 pairing (**Fig. 1c, Fig. S5**). Notably, bovine UBA7 can recruit mouse UBE2L6 to form disulfide complexes, and vice versa, but UBA7 from both species shows low affinity for human UBE2L6, likely due to species-specific barriers.

The enhanced formation of disulfide complexes in the context of UBA7 can be attributed to several critical factors. First, the exceptionally strong association affinity (1.2 µM) and specificity between UBA7 and UBE2L6 contribute to UBE2L6 being retained within the complex. This robust interaction effectively sequesters UBE2L6, preventing it from participating in downstream reactions. Unlike UBA1, which serves as the primary E1 enzyme for Ub and must interact with distinct E2 enzymes in the ubiquitylation cascade, UBA7 exclusively associates with UBE2L6 to conjugate ISG15. This specificity allows UBA7 to form a particularly tight and stable complex with UBE2L6. Specific and tight E1•E2 complexes have been characterized for other Ubl systems, including SUMO E1•Ubc9, UBA5•UFC1, and Atg7•Atg3, with respective K_D_ values of 1.26, 1.0, and 0.35 μM, respectively^29–32^.

Second, the conformational differences in the catalytic CCL of UBA7 compared to that of UBA1 play a pivotal role in facilitating or hindering the formation of the E1•E2 disulfide complex. Our cryo-EM structural analysis revealed that the CCL of UBA7 is shorter than that of UBA1, positioning it further away from the catalytic cysteine. This structural distinction results in UBA7 having a more exposed catalytic pocket. In line with these findings, our experimental manipulations of the CCL in both human UBA1 and bovine UBA7 confirmed the intrinsic importance of the CCL in modulating E1•E2 disulfide complex formation. Shortening the CCL of human UBA1 elicited an increased propensity for E1•E2 complex formation, indicating that the CCL influences the exposure of the catalytic cysteine for disulfide bonding. This observation is consistent with the determined crystal structures of UBA1, wherein the versatile CCL switches between a protective and a released flexible confirmation (**Fig. 4g**), and our HDX analysis (**Fig. 4e-f**).

Third, the pKa values of the active site cysteines in both E1 and E2 enzymes exert a substantial impact on their ability to form disulfide-bonded complexes. Lower pKa values imply a larger population of thiolate groups, making the cysteines more reactive for disulfide bond formation. Our measurements reveal that the pKa values of the catalytic cysteines in UBA7 (pKa 7.0) and UBE2L6 (pKa 8.6-9.0) are lower than that of UBA1 (pKa 7.5) and other E2 enzymes, which often have elevated pKa values exceeding 10^26^, rendering them more susceptible to disulfide bond formation. These lower pKa values likely arise from the unique demands of ISG15 conjugation, necessitating more rapid and efficient formation of the E1•E2 disulfide complex.

The phenomenon of disulfide bond formation involving active site cysteines in E1 and E2 enzymes has been implicated in other cellular contexts. For instance, the E2 enzyme Rad6 has been shown to form disulfide complexes in the presence of oxidants such as hydrogen peroxide (H_2_O_2_)^11^. Similarly, SUMO E1 and E2 form a disulfide complex under conditions of elevated ROS, such as arising upon DNA damage^33^. This process is highly sensitive to the ratio of thiol to thiolate groups on the cysteine residues. Interestingly, our findings herein indicate that deleting the CCL (dCCL) results in a similar phenomenon as observed with wild-type UBA1 and UBE2L3 under H_2_O_2_-induced oxidation, leading to the facilitation of disulfide complex formation (**Fig. S8**). This discrepancy may be attributed to the role of the CCL in protecting the catalytic cysteine.

ISGylation is an important immune response stimulated by interferon as an anti-microbial defense strategy^34–37^. Numerous proteins have been identified as hosting ISG15 modifications^20^. Our findings herein have broad implications that extend beyond a basic understanding of ISGylation. Research over the past two decades has revealed attenuation of SUMO’s E1•E2 complexes via direct oxidation or indirect ROS regulation^13,14,38^ resulted in a disequilibrium in the SUMO-conjugation cycle. Accordingly, we postulate that the rapid formation of the E1•E2 disulfide complex may represent a quiescent state upon UBA7 encountering UBE2L6. Under normal cellular conditions, this complex could serve as a protective mechanism, restraining an unnecessary ISGylation reaction cascade. However, during pathogenic infections or immune activation, such as through the STAT1/2 pathway, gene expression of ISG15-associated E1, E2, and E3 enzymes is triggered^36^. Similarly, the dynamic interplay between the UBA7•UBE2L6 complex and unbound forms of these proteins may function as a regulatory switch, finely balancing immune responses and cellular homeostasis (**Fig. 7a, b**).

**Figure 7.**
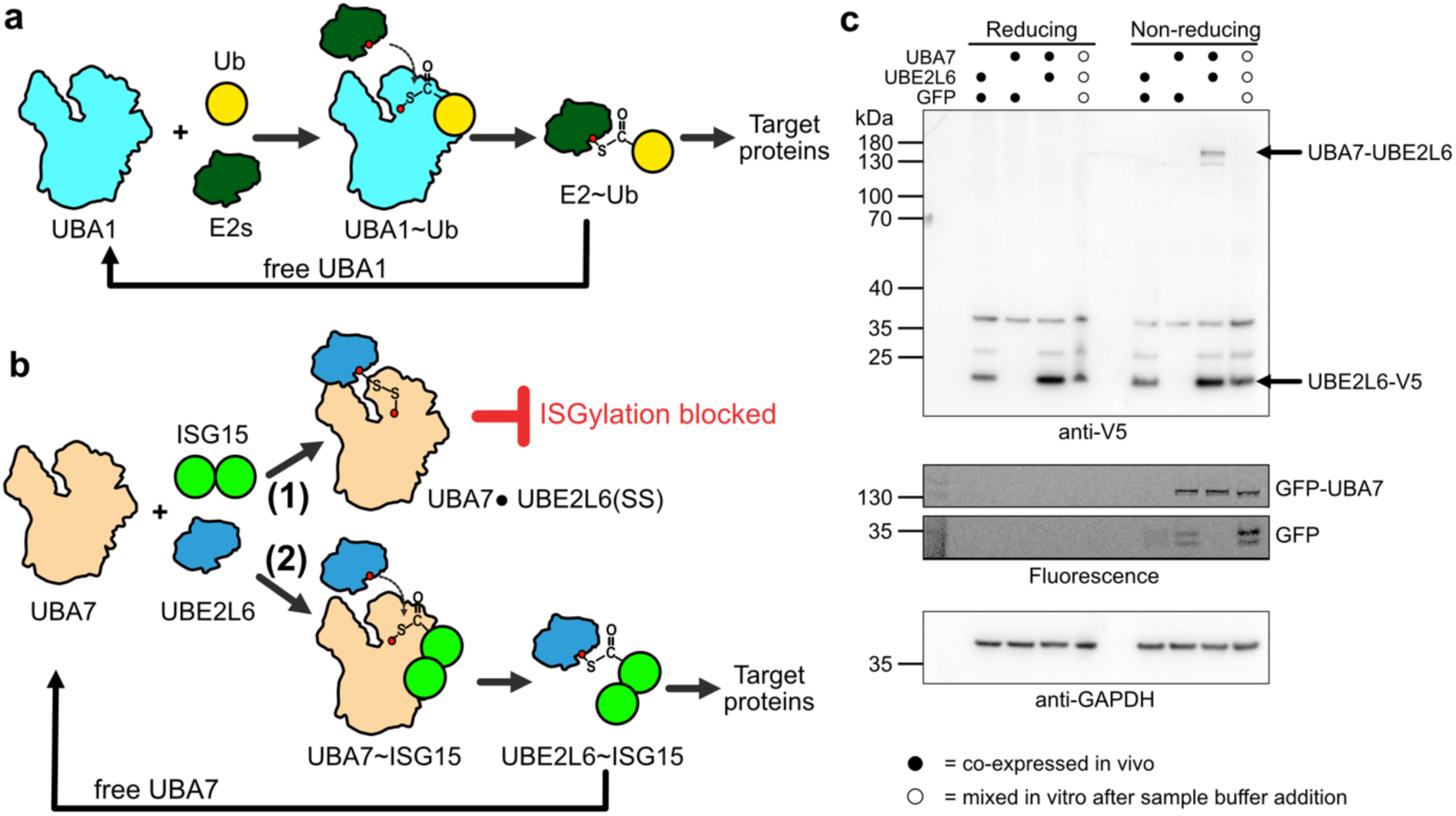
Summary of E1-E2 interactions in the Ub/Ubl cascade. **a.** In the ubiquitin reaction cascade, Ub is first charged by UBA1, followed by transfer from UBA1 to an E2 enzyme and sequential ubiquitylation reactions. When Ub is ultimately released, UBA1 is free to reinitiate the cascade. **b.** The ISG15 system operates via two distinct pathways. At the immune resting state, UBA7 and UBE2L6 favorably form a disulfide complex to suppress intracellular ISGylation (pathway 1). However, when the immune response is elevated, UBA7 charges ISG15 to initialize ISGylation of target proteins (pathway 2). **c.** Co-expressing GFP-UBA7 and UBE2L6-V5 in HEK293T cells. Non-reducing immunoblot of UBE2L6-V5 reveals a high molecular weight complex (>130 kDa) which is reduced by β-ME. The control groups (GFP and UBE2L6-V5, GFP and GFP-UBA7) demonstrate that the complex only forms when UBE2L6-V5 and GFP-UBA7 are co-transfected. Blue circles indicate the groups expressing UBE2L6 and UBA7 separately, then mixed after sample buffer addition. The fluorescent gel confirms GFP-UBA7 expression and anti-GAPDH is used as a loading control.

We then investigated whether the disulfide complex also occurs inside the cells (**Fig. 7c**). We co-expressed bUBA7 and bUBE2L6, tagged with GFP and V5, respectively, in HEK293T cells.

Consistently, we observed the formation of a UBE2L6-containing complex with the same molecular weight as UBA7•UBE2L6. This complex is sensitive to reducing agents such as β-ME. Importantly, we demonstrated that the complex can spontaneously form without any oxidant treatment. To confirm this, we separately expressed only UBA7 or UBE2L6 and mixed the samples after adding the sample buffer followed by immediate protein gel electrophoresis, ensuring that the complex was truly formed in the cell and not affected by the in vitro environment. The results support the findings in vitro. However, further comprehensive investigations are essential to unravel ISG15-specific cellular responses in terms of oxidation-reduction dynamics and the UBA7•UBE2L6 disulfide complex.

In conclusion, our study sheds light on the complicated and specialized mechanisms governing ISG15 conjugation through the formation of a covalent disulfide complex between UBA7 and UBE2L6. The specificity, structural nuances, and redox sensitivity of this interaction provide valuable insights into the broader landscape of ubiquitin-like modifications and how they are regulated. Further investigations into the role of UBA7 in redox regulation and its implications in immune responses promise to unveil novel avenues for understanding cellular defense mechanisms and the development of therapeutic interventions.

## Materials and Methods

### Protein Expression and Purification

Custom gene synthesis was employed for all wild-type genes, except UBA1 and human UBA7. The human UBA1 and UBA7 genes were obtained from Addgene (34965, 12438). Subsequently, all genes were subcloned into the pRSFDuet-1 vector (Novagen), with the addition of a polyhistidine affinity tag (His-tag: His6, His8-GST, or His8-MBP) featuring a TEV protease cleavage sequence. Chimeric versions of UBA1 or UBA7 were constructed through a series of multi-step polymerase chain reactions. Single or multi-point mutations were introduced into E1 and E2 enzymes or Ubl through site-directed mutagenesis. A comprehensive list of genes and proteins utilized in this study is presented in Table S1. Each plasmid was individually transformed into the *Escherichia coli* BL21 RIL cell line (Agilent) to initiate protein overexpression. Protein expression was induced by adding 0.6 mM ITPG when cell cultures reached an optical density of 0.6 to 0.8. UBA1 and UBA7 were expressed at 16 °C, whereas E2 enzymes, ISG15, and Ub were expressed at 20-25 °C. Human UBA7 was purchased directly from R&D Biosystems (catalog #: E-307).

The purification process began by purifying the His-tagged proteins through gravity open columns using cOmplete resin (Roche, USA), before subjecting them to concurrent dialysis and proteolysis. Cleaved products were subsequently reloaded onto open columns and the flow-through and wash fractions were collected. E1 and E2 enzymes underwent further purification via ion-exchange chromatography, followed by size-exclusion chromatography utilizing a HiLoad Superdex200 16/60 column (Cytiva, USA). ISG15 and Ub were directly processed through a Superdex 200 or Superdex 75 10/300 increase column, respectively. Eluted fractions were assessed via SDS-PAGE gels, and pure fractions were combined, concentrated, aliquoted, and promptly frozen using liquid nitrogen. The proteins were then stored at -80 °C.

### SEC-MALS Analysis

To determine the molecular weights of UBA7, UBE2L6, and their complex, we employed Size Exclusion Chromatography coupled with Multi-Angle Light Scattering (SEC-MALS). Samples containing 1 mg/ml of UBA7 or UBE2L6, as well as a mixture of UBA7 and UBE2L6 (at a 1:5 molar ratio), were prepared separately for SEC-MALS analysis. Then, 100 µl of each sample was injected into a Superdex 200 10/300 increase column integrated into an Agilent 1260 Infinity HPLC system. Detection was performed using a minDAWN TREOS detector equipped with three angles (43.6°, 90°, and 136.4°) and a 659 nm laser beam. To analyze the sizes of individual proteins and complexes, an Optilab T-rEX differential refractive index detector (Wyatt Technology, USA) was connected to the apparatus. Calibration of the detectors was carried out using bovine serum albumin (BSA) as a reference. Molecular weights of UBA7, UBE2L6, and disulfide-bonded UBA7•UBE2L6 complex were computed using ASTRA6 software (Wyatt Technology, USA), with a refractive index increment (dn/dc) value of 0.185 ml/g. The solvent’s refractive index was defined as 1.331, with a viscosity of 0.894 cP.

### Octet Bio-Layer Interferometry

A ForteBio Octet96 system (Sartorius) was utilized to determine the binding equilibrium constants between various E1 and E2 enzyme pairs. We purified and concentrated His-GST-tagged E1 variants, including wt UBA7, UBA7(C597A), UBA7(eCCL), UBA7(U1), UBA1, and the UFD domain of UBA7 alone. These concentrated proteins were diluted into Octet running buffer (comprising 25 mM HEPES pH 7.5, 150 mM NaCl, 0.01% Tween 20, and 0.1 mg/mL BSA). Kinetic assays were initiated by capturing the GST-fused E1 variants. Anti-GST biosensors were used to load approximately 10-50 μg/ml of these variants. After washing away unbound E1 variants, the E1-captured biosensors were immersed in wells containing E2 proteins including UBE2L3(C86A), UBE2L6(C86A), and wt UBE2L6. All binding kinetics experiments were conducted in triplicate. Steady-state responses (measured in nanometers) corresponding to various E2 protein concentrations were extracted for fitting using a single-phase 1:1 binding model with the Octet data analysis software.

### Hydrogen-Deuterium Exchange Mass Spectrometry (HDX-MS)

HDX-MS experiments were conducted employing an automated HDX robot (LEAP Technologies, USA) coupled with an M-Class Acquity LC and HDX manager (Waters, USA). A 3 μL protein solution containing 80 μM of protein samples in equilibration buffer (25 mM Tris 200 mM NaCl buffer, pH 7.6) was combined with 57 μL of labeling buffer (25 mM Tris 200 mM NaCl buffer, pD 7.6) and incubated at 4 °C for 30, 60, 600, 1800, 3600, or 14400 seconds. After the labeling reaction, quenching was achieved by adding 50 μL of the labeled solution to 50 μL of quench buffer (25 mM Tris 200 mM NaCl buffer, pH 2.2), resulting in a final quench pH of ∼2.5. Subsequently, 50 μL of the quenched sample was passed through an Enzymate BEH Pepsin column (Waters) at a flow rate of 70 μL/min (15 °C) and a VanGuard Pre-column Acquity UPLC BEH C18 (Waters) for 3 minutes. The pepsin-digested peptides were then transferred to a C18 column and separated using a gradient from 5% to 40% solvent B (acetonitrile with 0.2% formic acid) at 4 °C, with solvent A being 0.2% formic acid in water. Mass spectrometric acquisitions were conducted in positive and sensitivity modes over the m/z range of 50–2000 Da using a Synapt G2 HDMS mass spectrometer (Waters) with a standard electrospray ionization source. All samples were independently analyzed three times for data repeatability. Peptides were identified through MSE analysis, and intermittent infusion of (Glu1)-fibrinopeptide B human (CAS No 103213-49-6, Sigma-Aldrich) was employed for lock mass correction, with an expected m/z value of 785.8426. HDX data were subsequently analyzed using PLGS (v3.0.2) and DynamX (v3.0.0) software provided with the mass spectrometer. Peptide identification in DynamX was guided by the following criteria: minimum intensity of 1000, maximum ppm error of 25, and a file threshold of 3/3.

### *In vitro* Biochemical Assays of Enzymatic Activities

To enable crosslinking with fluorescein maleimide (AnaSpec, USA), a cysteine residue was introduced at the N-terminus of Ubl proteins, according to our previously published protocol^39,40^. The assays were conducted in a buffer containing 25 mM HEPES pH 7.5 and 200 mM NaCl.

Samples were quenched using 4X non-reducing Laemmli SDS sample buffer (TPpro, Taiwan) and then separated on 4-20% gradient SDS gels (GenScript, USA). Visualization of protein bands was carried out using an iBright FL1000 system (ThermoFisher, USA) with visible light or the default Alexa488 fluorophore profile (excitation at 488 nm, emission at 530 nm). Protein band signals were analyzed using open-source Fiji software (https://github.com/fiji/fiji). To prevent multiple crosslinking reactions, native cysteine residues in Ubl proteins were mutated to alanine (denoted as ISG15* and Ub*, with the asterisk signifying fluorescein).

Charging of Ubl protein (ISG15* or Ub*) onto the corresponding E1 enzyme was accomplished by mixing 1 µM E1 enzyme, 5 µM Ubl, 5 mM ATP, and 5 mM MgCl_2_ in the assay buffer. The reaction proceeded at 37 °C for 30 minutes. Transthioesterification reactions of Ubl proteins to E2 enzyme were carried out by incubating E2 enzyme and E1∼Ubl* at 37 °C for 30 minutes, followed by immediate quenching for SDS-PAGE analysis.

Kinetic experiments related to E2∼Ubl* thioester formation involved mixing 0.3 µM E1∼ISG15* (UBA7 or its variants) and 0.5-8 µM UBE2L6 (8-64 µM UBE2L6 for UBA7(U1)) on ice for 20 seconds, followed by direct quenching. The UBE2L6∼ISG15* fluorescence signals were detected in the SDS gels for kinetic analysis. Disulfide-bonded E1•E2 complexes were formed by mixing 0.5 µM E1 enzyme (UBA7 or UBA1 variants) and varying concentrations of E2 enzyme (UBE2L3 or UBE2L6) in the assay buffer. These reactions were incubated at 25 °C using a thermal cycler for 5-10 minutes and then stopped using 4X non-reducing Laemmli buffer. The E1•E2 complexes in the SDS gels were visualized by Coomassie blue staining for kinetic analysis.

### Determination of Thiol pKa values for Catalytic Cysteines in E1 and E2 Enzymes

To determine the pKa values of E2 enzymes, all non-catalytic cysteines in E2 were mutated to serine or E2 enzymes with only one cysteine were generated by total gene synthesis to ensure that the pKa values were solely associated with the catalytic cysteine. C-terminal His-tagged E2 enzyme was then purified and concentrated as described above. Isothermal titration calorimetry (ITC) was employed to detect heat changes upon reaction of cysteine thiols to iodoacetamide^25,26^. Alkylation is pH-dependent and irreversible, making it a suitable indicator for directly measuring cysteine thiol pKa values. To prepare the E2 enzyme samples, their pH was adjusted as necessary using a MidiTrap G25 column. The sample buffer contained 25 mM HEPES, CHES, or CAPS, covering a pH range from 7 to 10.5, with the final protein concentration set at 80 µM. The experiments were conducted using a Microcal iTC200 system (Malvern Panalytical), involving a single injection of 2 µl of 80 mM iodoacetamide into 280 µl of buffer-exchanged E2 enzyme at 25 °C, with the paddle stirring at 750 rpm. Endothermic thermograms (dQ/dt) were generated. E2-free buffer or iodoacetamide-free injection samples were used as experimental controls to monitor heat changes over time. The dQ/dT data were subjected to iterative non-linear regression based on the Henderson-Hasselbach equation^25,26^ to obtain pKa values for the catalytic cysteines of E2 enzymes.

Since both UBA1 and UBA7 possess too many cysteines to enable mutation-based pKa measurements by means of ITC, we monitored E1∼Ubl thiolation yields in buffers of varying pH (from 6 to 9.5), both with and without iodoacetamide treatments. We anticipated the catalytic cysteine would be fully alkylated at low pH. However, both E1 enzymes showed instability at acidic pH levels (5.5 or lower) due to rapid precipitation. To address this issue, the E1 enzymes (10 µM) were buffer-exchanged into the desired pH buffer and reacted with 1 mM iodoacetamide for 5 minutes at 25 °C to block some of their catalytic cysteines. Aliquots of 2 µl were sampled from the incubation mixture and quickly added to 18 µl of assay buffer at pH 7.5, containing 5 µM Ubl*, 5 mM ATP, and 5 mM MgCl_2_ for Ubl* charging experiments. Iodoacetamide-free buffers were used as internal controls to quantify the percentages of E1∼Ubl* on SDS gels according to fluorescence signal. The pH-dependent yields of E1∼Ubl* were plotted and fitted using the same equation deployed for E2 enzymes to determine the pKa values of the catalytic cysteine of E1 enzymes.

### Cryogenic Electron Microscopy (Cryo-EM) data collection, analysis and model building

For the MBP-fused UBA7•UBE2L6 disulfide complex, we incubated a mixture of 20 µM purified MBP-UBA7 and 100 µM UBE2L6 for 30 minutes at 37 °C without additional linker molecules, and excess UBE2L6 was removed by size-exclusion chromatography to obtain the purified disulfide complex. The resulting complex (0.5 mg/mL) was applied to Quantifoil Cu grids (300 mesh, R1.2/1.3 µm) using a Vitrobot Mark IV system at 4 °C and 100% humidity. Blotting force and time were set to 0 and 3-3.5 seconds, respectively. Grids were subsequently flash-frozen in liquid ethane and stored in a liquid nitrogen reservoir. The data collection was carried out on a Titan Krios microscope equipped with a K3 direct detector (ThermoFisher, USA). A total of 10548 50-frame movies were collected in super-resolution pixel resolution of 0.324 Å, with a total electron dose of 50 e-/Å^2^. Data processing was performed using cryoSPARC 4.5, following the standard protocol for single-particle reconstruction. The process included movie alignment using Patch motion, CTF estimation using PatchCTF, blob particle picking, *ab initio* model building, heterogeneous refinement, homogeneous refinement and, finally, selection of 46,385 particles to reconstruct the MBP-UBA7•UBE2L6 complex (**Fig. S2**).

For the activation complex, 6.5 µM UBA7•UBE2L6 was mixed with 30 µM of ISG15 in the presence of 5 mM ATP and 5 mM MgCl_2_. The reaction was carried out at 25 °C for 20 minutes before vitrification. 4.5 µM of UBA7•UBE2L6∼ISG15(A) was applied to Quantifoil grid followed by a vitrification process as mentioned in this session. A total of 18731 micrographs were collected from 3 separate acquisitions. The acquisition parameters and process procedures were performed following the established protocol used in the MBP•UBA7-UBE2L6 case. In short, three ab initio models were used followed by multiple filtrations of heterogeneous refinements. 176,617 particles were used for non-uniform refinement resulting in a 2.99 Å map evaluated by the gold standard FSC_0.143_ criterion. The two half-maps were further post-processed using DeepEMhancer^41^ for model building and map deposition (**Fig. S3**). The structure was built using the AlphaFold3 template of UBA7-UBE2L6-ISG15. Sidechain rotamers, backbone traces, and AMP molecules were interactively modeled and inspected using Phenix Real-space refinement^42^ and Coot^43^. The final model was evaluated with MolProbity and the Phenix validation tool to ensure publication quality and suitability for further analysis.

### Detection of covalent disulfide bonding between bovine UBA7 and UBE2L6 in cells

For the detection of covalent bonding, the cell extracted from 293T cells that transiently expressed GFP-UBA7 (bovine), UBE2L6-V5 (bovine) separately, or both genes at the same time. The cells were lysated by 1X Laemmli buffer (50 mM Tris-HCl, pH 6.8, 10% glycerol, and 2% SDS) with or without the addition of 100 mM β-ME. To confirm the covalent disulfide UBA7•UBE2L6 formation, cell extracts containing GFP-UBA7 and UBE2L6-V5 separately were mixed for 10 minutes at room temperature prior to the electrophoresis. Immunoblotting was then performed as described previously^44^. The V5 antibody (AB3792, 5,000x dilution) was from Sigma-Aldrich; the antibody against GAPDH (GTX627408, 12,000x dilution) was from GeneTex. Anti-mouse IgG (JK715-035-150) and anti-rabbit IgG (JK711-035-152) were from Jackson ImmunoResearch.

## Acknowledgments

We appreciate the support provided by the Academia Sinica (AS) (Taipei, Taiwan) Career Development Award (AS-CDA-110-L03) and from the National Science and Technology Council (110-2311-B-001-046 and 111-2113-M-001-026-MY2) to K.-P.W.. We are also thankful for the technical guidance and instrumental resources made available to us by Dr. Shu-Juan Jao in the AS Biophysics Core Facility (AS-CFII-111-201), Dr. Shu-Yu Lin in the AS Common Mass Spectrometry Facilities (AS-CFII-111-209), Eric Yen in the AS Grid Computing Center (AS-CFII-112-103), Dr. Chun-Hsiung Wang and Dr. Yuan-Chih Chang in the AS Cryo-EM Center (AS-CFII-112-210), and Dr. Meng-Ru Ho in the Biophysics Instrumentation Laboratory, Institute of Biological Chemistry, AS.

## Supplementary Figures

**Supplementary Figure 1.**
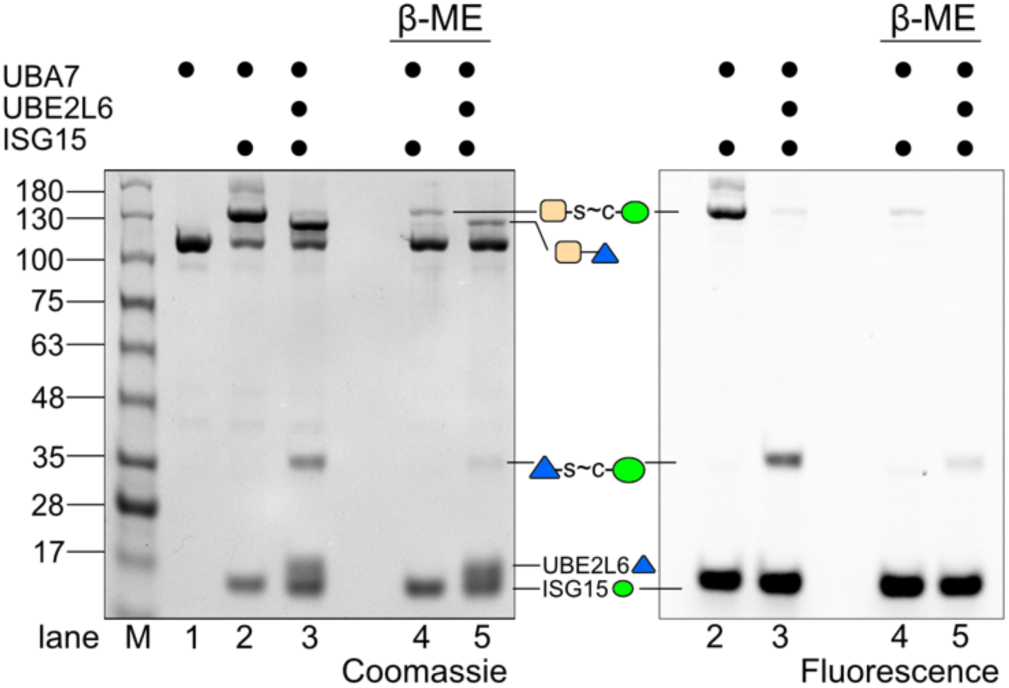
The UBA7•UBE2L6 complex is sensitive to a reducing agent. UBA7 was used to charge ISG15 and to transfer it from UBA7 to UBE2L6. After reaction at 37 °C for 30 mins, the reaction was quenched using 4X non-reducing Laemmli sample buffer. β-mercaptoethanol (β-ME, 10 mM) was also added in the experimental (reduced) samples. The SDS gel shows that the thioester-linked products, including UBA7∼ISG15* and UBE2L6∼ISG15*, are disrupted by β-ME treatment, as observed in both Coomassie-stained and fluorescence images. A band corresponding to the UBA7•UBE2L6 complex is only detected in the Coomassie-stained gel, and it is also sensitive to β-ME treatment, implying a disulfide-bonded complex.

**Supplementary Figure 2.**
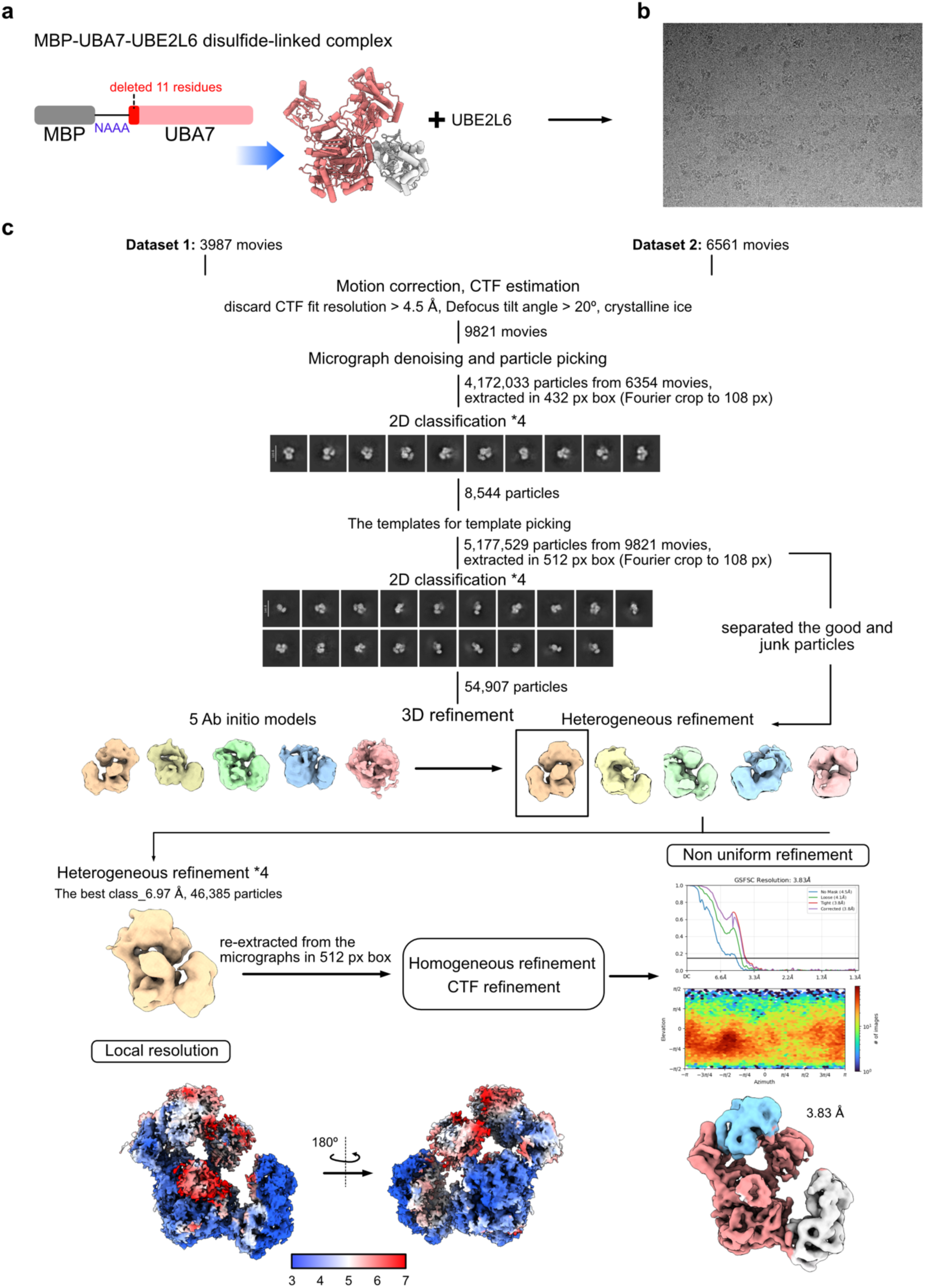
Cryo-EM analysis of the MBP-UBA7•UBE2L6 complex (UBA7•UBE2L6) **a.** Schematic representation of the formation of the MBP-UBA7•UBE2L6 complex. The first 11 residues of UBA7 have been deleted and replaced by a linker sequence (-NAAA) attached to the C-terminus of maltose-binding protein (MBP). **b.** The representative micrograph out of all images used for processing. **c.** The cryo-EM data processing flow chart. The final map was reconstructed to 3.83 Å.

**Supplementary Figure 3.**
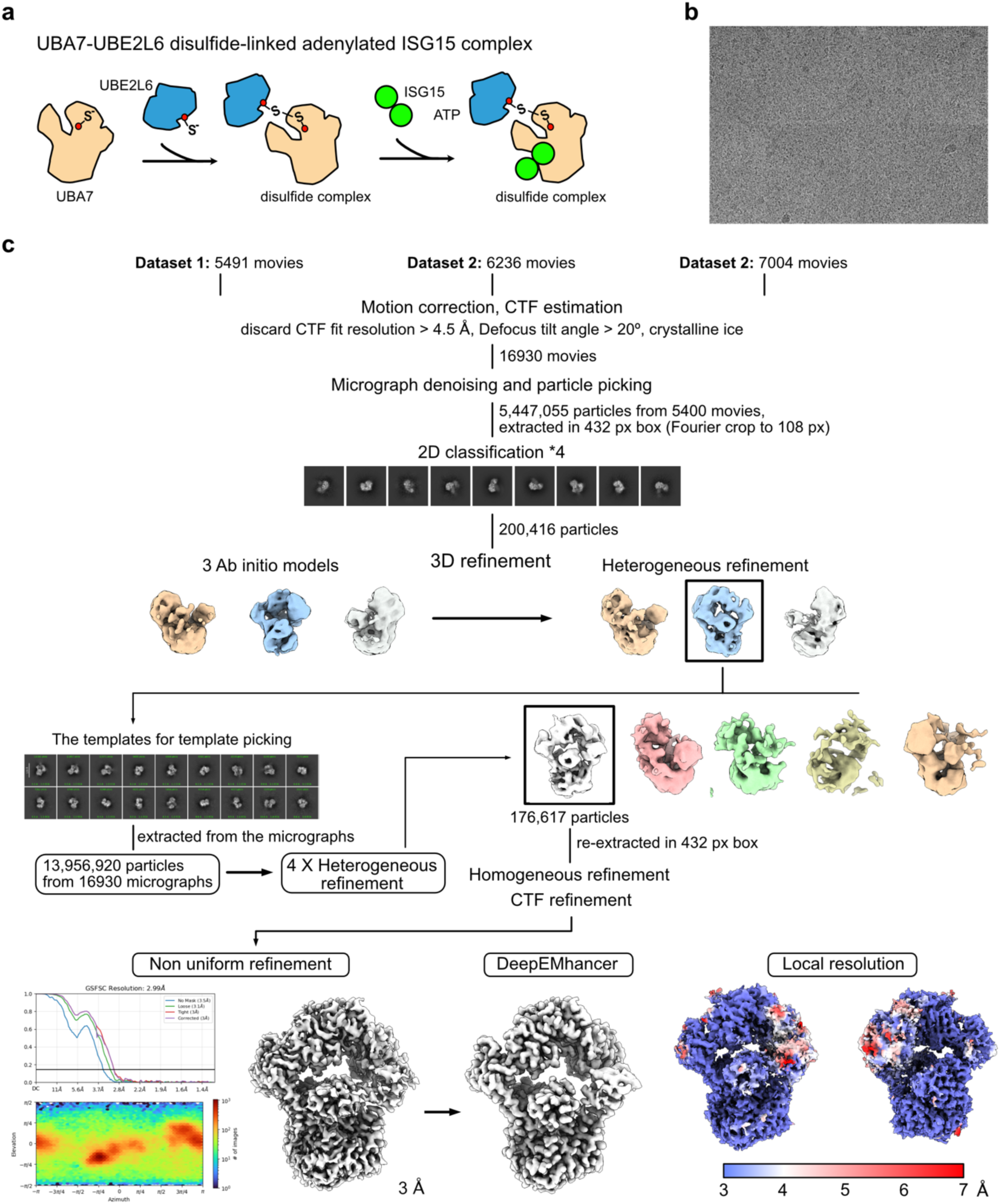
Cryo-EM data processing of the UBA7•UBE2L6 adenylated ISG15 complex (UBA7•UBE2L6-ISG15(a)) **a.** Schematic depicting the formation of the UBA7•UBE2L6-ISG15(a) complex. In the complex, the active site cysteines of UBA7 and UBE2L6 are cross-linked by a disulfide bond. **b.** The representative micrograph out of all images used for processing. **c.** The cryo-EM data processing flow chart. The final map was reconstructed to 2.99 Å whereas local resolution was estimated (FSC_0.143_) suggesting FCCH and SCCH domains exhibit lower resolution (i.e. 4-7 Å).

**Supplementary Figure 4.**
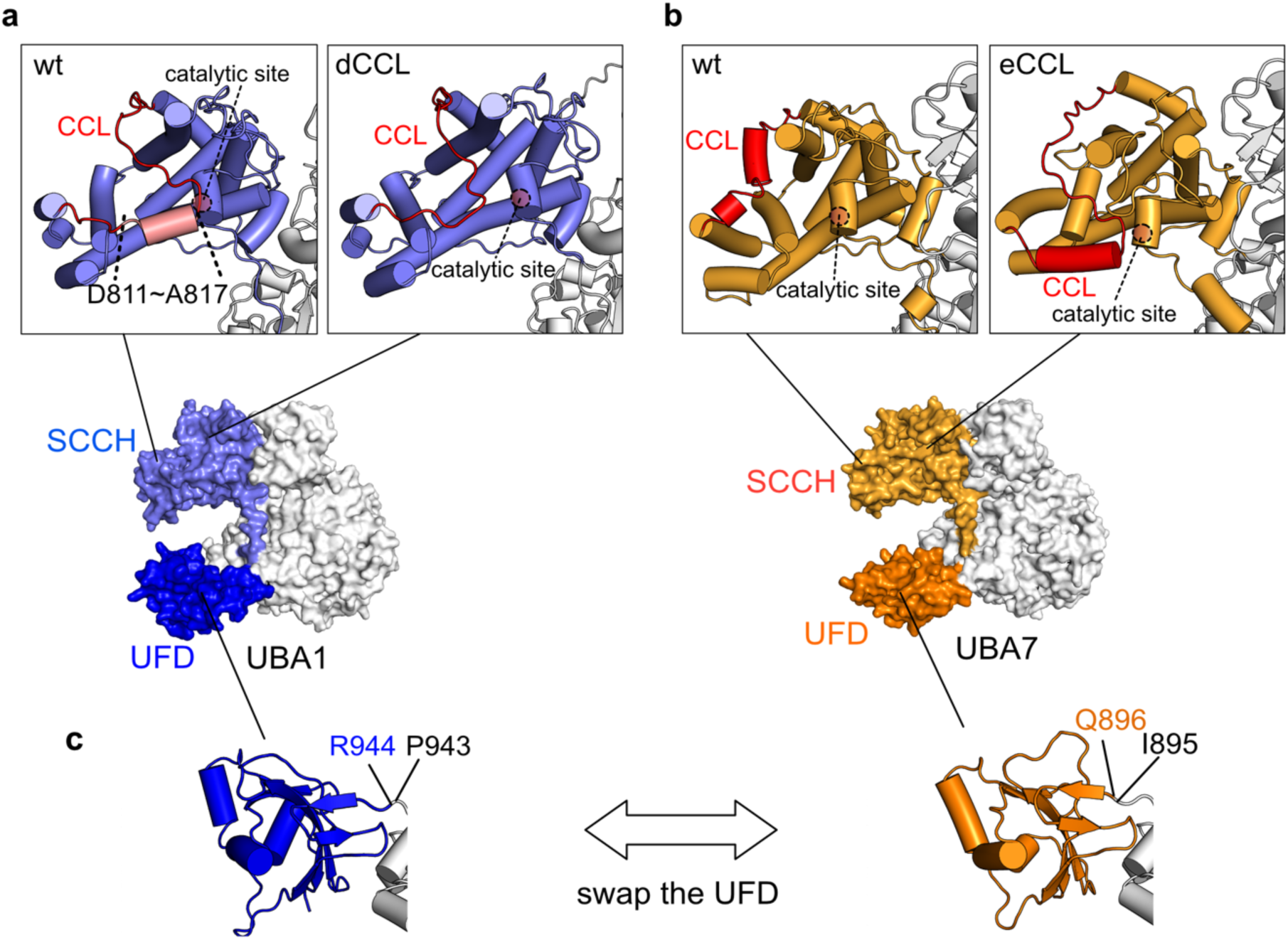
Models of chimeric proteins for functional analyses of E1 enzymes. **a-b.** Extension or deletion of the cysteine capping loop (CCL) region in UBA7 and UBA1, respectively, results in the catalytic cysteine either being exposed or covered. **c.** In the case of the chimeric E1 variants UBA7(U1) and UBA1(U7), UFD domains were swapped while their linker regions remained unchanged. Substitutions of the UFDs were initiated at amino acid positions Q896 and R944 for UBA7 and UBA1, respectively. These positions correspond to residues located between the first β-sheet of the UFD and the linker.

**Supplementary Figure 5.**
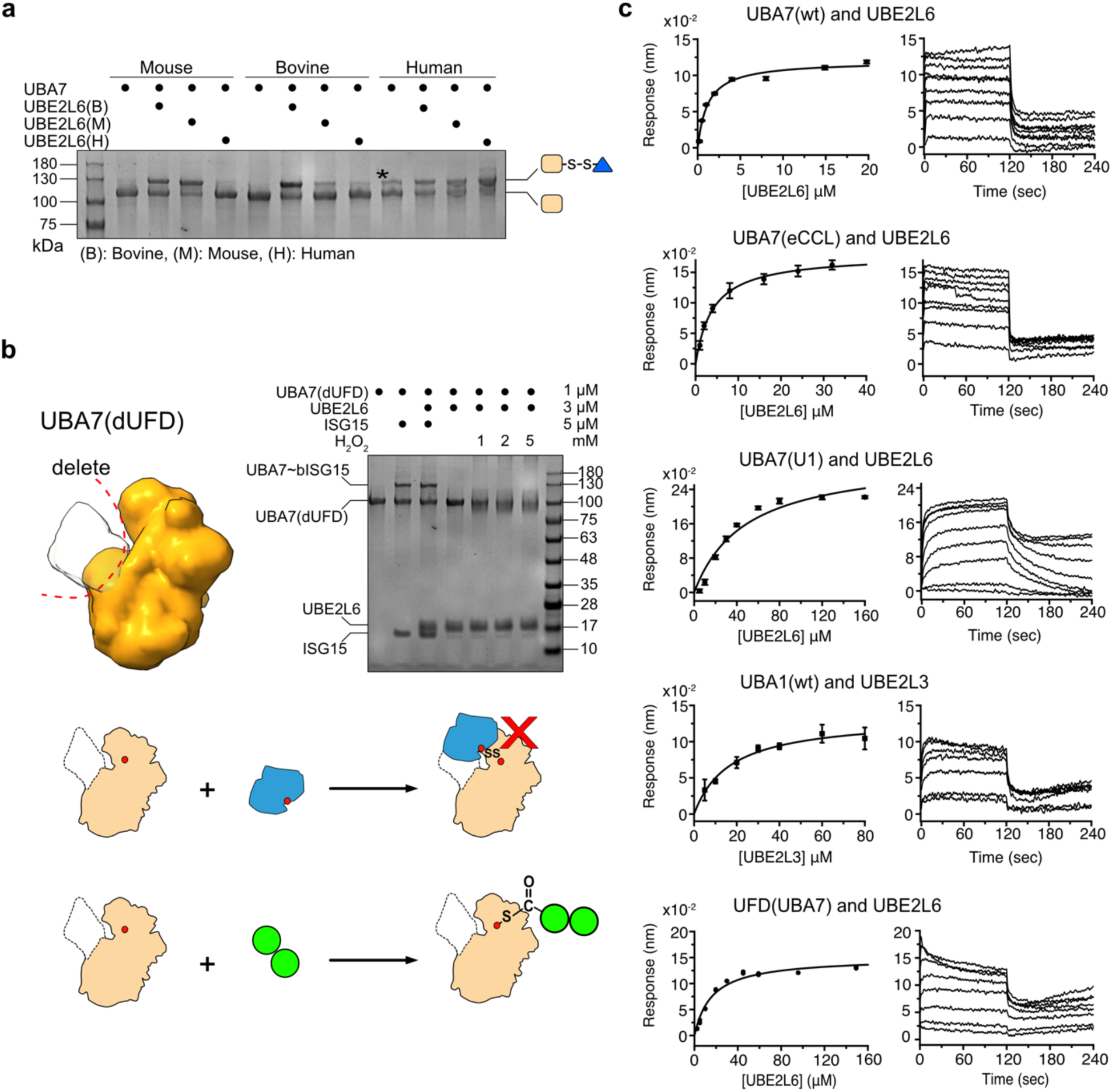
a. Disulfide bond formation assay of UBA7 and UBE2L6 among three species: mouse (M), bovine (B), and human (H). UBA7 and UBE2L6 were mixed and incubated for 30 minutes at 37 °C, resulting in cross-reactivities. A contamination band for hUBA7 (R&D Biosystems, USA) is indicated by *. **b.** UFD-deleted UBA7 was used to validate ISG15 activation, transthioesterification, and UBA7(dUFD)•UBE2L6 complex formation. Use of UBA7(dUFD) does not impair ISG15 activation, but transthioesterification and disulfide complex formation are not observable in the Coomassie-stained SDS gel. H_2_O_2_ treatment of an UBA7(dUFD) and UBE2L6 mixture results in no disulfide complex formation. **c.** Curve fitting and sensorgrams of BLI-determined binding affinities for E1-E2 pairs. The UBA7(C597A) or UBE2L6(C86A) mutant was used for BLI experiments on wtUBA7•UBE2L6 or UBA7(eCCL) •UBE2L6, respectively, to avoid irreversible disulfide complex formation during the BLI experiments.

**Supplementary Figure 6.**
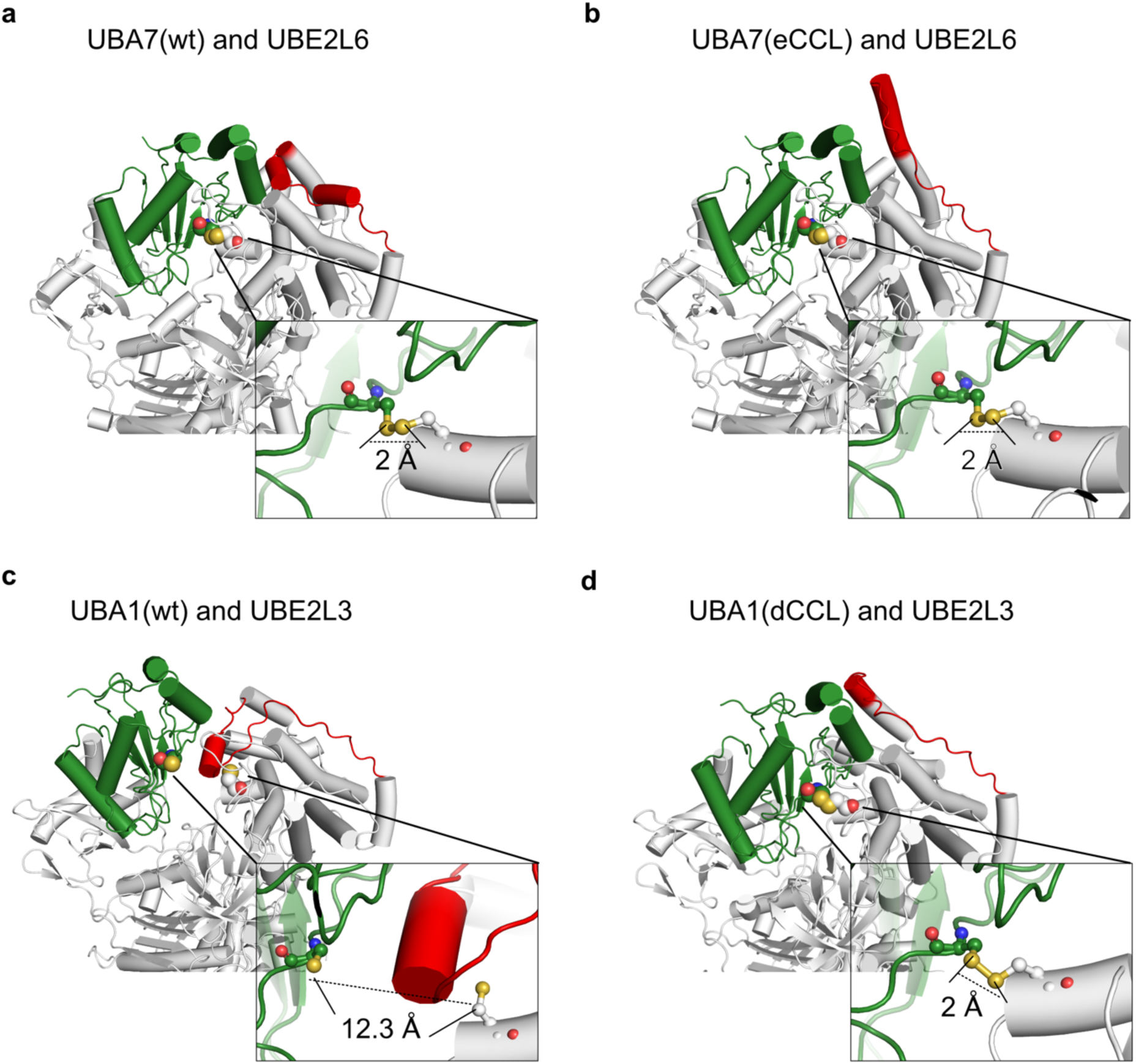
AlphaFold2-generated models of complexes comprising E1 variants and their cognate E2 enzymes. AlphaFold2-predicted E1•E2 complex structures, including wt UBA7•UBE2L6 (**a**), UBA7(eCCL)•UBE2L6 (**b**), wt UBA1•UBE2L3 (**c**), and UBEA1(dCCL)•UBE2L3 (**d**). E1, E2 and the CCL are colored in gray, green and red, respectively. Zoomed-in views of the contacts near the two active site cysteines show the distances between the two sulfur atoms of the cysteines. Both wt and eCCL variants of UBA7 can form disulfide complex with UBE2L6, but only the UBA1(dCCL) variant can do so.

**Supplementary Figure 7.**
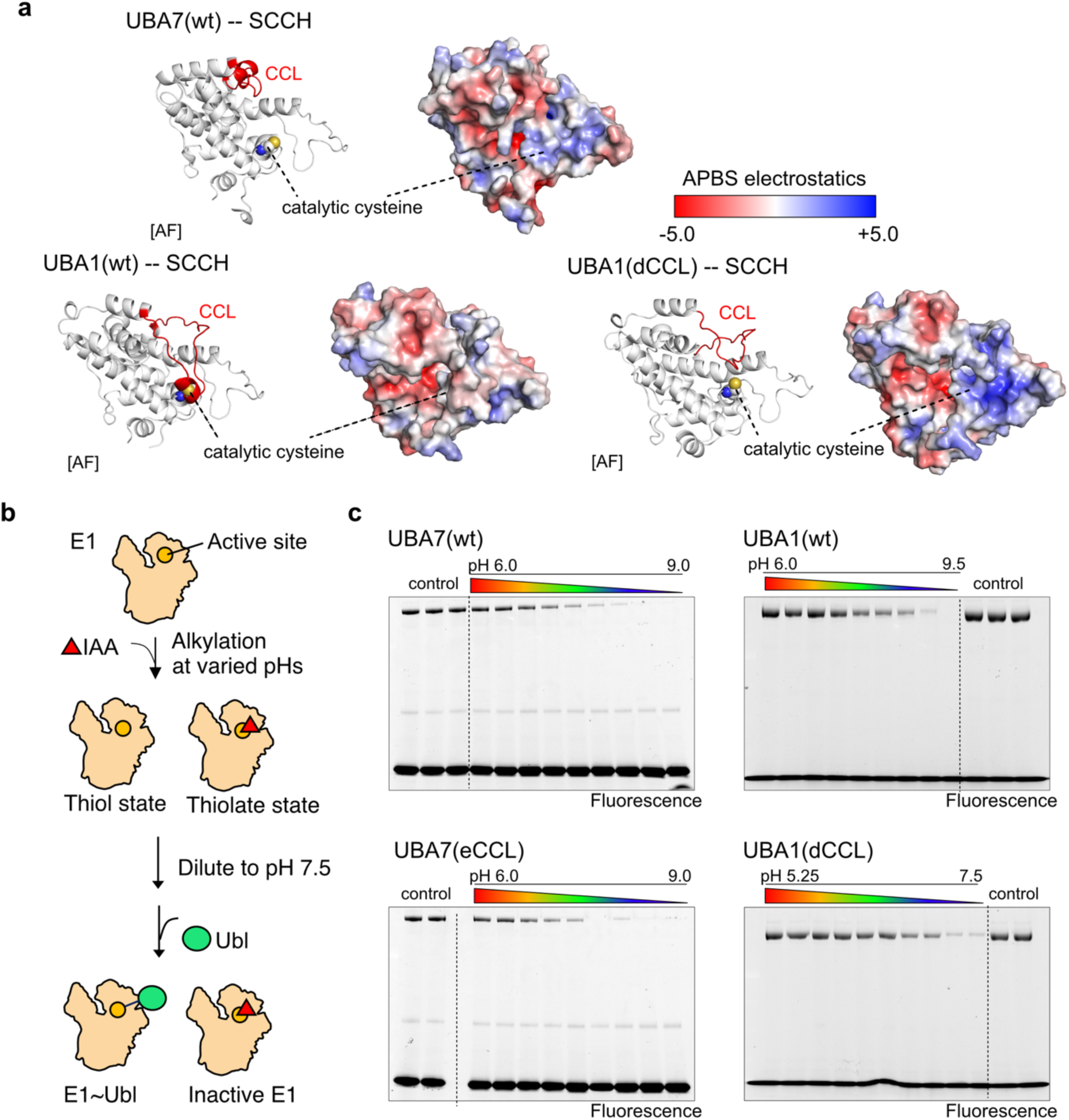
pH-dependent inactivation of the active site cysteine of E1 enzymes. **a.** The catalytic cysteine of E1 enzymes is influenced by the local acidic patch, as revealed by the electrostatic surface potential drawn in PyMOL 2.2. Cartoon and surface views show the locations of the CCL and catalytic cysteines. **b.** E1 enzymes were alkylated at desired pH, then diluted to pH 7.5, before undergoing an immediate E1∼Ubl charging reaction. **c**. At higher pH, greater amounts of alkylated E1 result in lower quantities of E1∼Ubl* product. The ratios of alkylated E1 versus pH were used to obtain the pKa values for each E1 enzyme. Control lanes are E1 enzymes without alkylation, eliciting maximal amounts of E1∼Ubl* product.

**Supplementary Figure 8.**
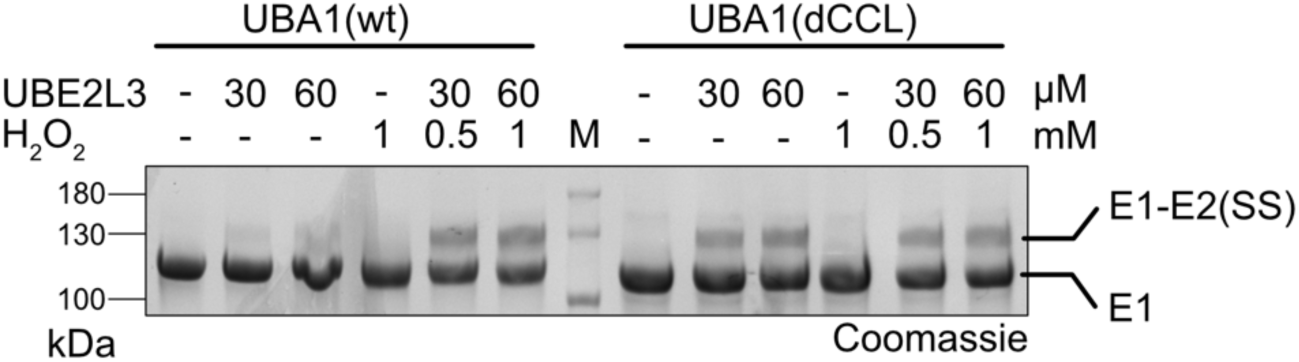
UBA1 forms disulfide complex with UBE2L3 in ROS solution. UBA1•UBE2L3 disulfide complex formation was tested in the presence of H_2_O_2_. In the absence of H_2_O_2_, wt UBA1 cannot form significant amounts of UBA1•UBE2L3 (SS) complex, even when a 40-fold concentration of UBE2L3 (60 µM) to UBEA1 (1.5 µM) was used. In contrast, upon addition of 0.5 mM H_2_O_2_, strong signal of disulfide complex was detected. When the CCL-shortened UBA1(dCCL) variant was used, disulfide complex spontaneously formed in the absence of H_2_O_2_, and H_2_O_2_ treatment did not further facilitate the ability of the UBA1(dCCL) variant to generate disulfide complex.

**Supplementary Table 1.**

| Gene name | Source | Mutation | Used for | Uniport ID |
| --- | --- | --- | --- | --- |
| hUBA1 | Addgene | no | HDX / General assay | P22314 |
| hUBA1(dCCL) | This study | yes | General assay | - |
| hUBA1(U7) | This study | yes | General assay | - |
| bUBA7 | Synthesized | no | General assay | Q5GF34 |
| EGFP-bUBA7 | This study | no | General assay | - |
| MBP-bUBA7 | This study | no | General assay | - |
| bUBA7(C597A) | This study | yes | HDX / General assay | - |
| bUBA7(eCCL) | This study | yes | General assay | - |
| bUBA7(U1) | This study | yes | General assay | - |
| bUBA7(dUFD) | This study | yes | General assay | - |
| hUBA7 | R&D Biosystems | no | General assay | P41226 |
| mUBA7 | Synthesized | no | General assay | Q9DBK7 |
| GST-bUBA7(C597A) | This study | yes | BLI assay | - |
| GST-bUBA7(eCCL) | This study | yes | BLI assay | - |
| GST-bUBA7(U1) | This study | yes | BLI assay | - |
| GST(bUFD) | This study | no | BLI assay | - |
| GST-hUBA1 | This study | no | BLI assay | - |
| UBE2L6 (Human) | Synthesized | no | General assay | O14933 |
| UBE2L3 (Human) | Synthesized | no | HDX / General assay | P68036 |
| UBE2L6 (Mouse) | Synthesized | no | General assay | Q9QZU9 |
| UBE2L6 (Bovine) | Synthesized | no | General assay | A5PJC4 |
| UbcH10-His<br>(single cysteine, C114) | Synthesized | yes | ITC (pH assay) | O00762 |
| UBE2L6-His<br>(bovine, single cysteine, C86) | Synthesized | yes | HDX / ITC / BLI<br>assay | - |
| UBE2L6-V5<br>(bovine, single cysteine, C86) | This study | yes | General assay | - |
| UBE2L3<br>(Human, single cysteine, C86) | Synthesized | yes | ITC (pH assay) | - |
| UBE2L6-His<br>(Human, single cysteine, C86) | Synthesized | yes | ITC (pH assay) | - |
| UBE2L(C86A) | This study | yes | BLI assay | - |
| ISG15 (Bovine) | Synthesized | yes | General assay | O02741 |
| Ub (Human) | Synthesized | yes | General assay | - |

